# Synteny and linkage decay in bacteriophage pangenomes

**DOI:** 10.1101/2025.08.12.669904

**Authors:** Jemma M. Fendley, Marco Molari, Richard A. Neher, Boris I. Shraiman

## Abstract

Bacteriophages are rich in genetic diversity, due to frequent horizontal transfer and recombination. This makes traditional microbial phylogenetic analyses, often based on the assumption of vertical inheritance, not suitable for interpreting this diversity. Here, inspired by recent work on bacterial pangenomes, we investigate the evolution of a collection of 3425 actinobacteriophage genomes. We find that synteny is strongly conserved: core genes have a well-defined order, and most accessory genes are localized in a few locations along the core genome backbone. Within the core genome alignment, linkage disequilibrium decays rapidly with distance in some groups, while phylogenetic structure in other groups causes long-range linkage. Our quantitative characterizations extend across many groups of phages and indicate widespread homologous recombination restricted by strong gene order conservation.

## 1 Introduction

Bacteriophages (phages), viruses which infect bacteria, are ubiquitous in nature and thought to be the most abundant biological entity on this planet. They play a crucial role in shaping microbial ecology and evolution: they facilitate gene transfer between bacteria [1], control population sizes [2], generate diversity [3], and drive bacterial evolution of anti-phage defense systems [4]. Understanding the evolution of phages and their bacterial hosts is crucial for human health, with vast applications from the gut microbiome [5] to phage therapy, a promising route in the fight against antimicrobial resistance [6].

Due to their high abundance, ancient evolutionary origins, and frequent horizontal gene transfer and recombination, phages are extremely genetically diverse. Unlike their bacterial hosts, phages have no universal gene, like 16S rRNA, upon which to build a phylogenetic tree [7], leading to historically dynamic taxonomic classifications [8]. Moreover, genetic exchange between distantly related strains has been demonstrated [9]. These features support the idea that phage genomes should be thought of as ‘mosaic’, a concept which views each phage genome as a set of interchangeable blocks [10, 11]. Mosaicism is well-established, with extensive evidence across different groups of phages [12, 13], exhibiting different rates [14].

Yet, not enough is systematically understood about the extent and breadth of this mosaicism. Can we characterize the processes that generate this mosaicism? What blocks are being interchanged, how often, and between whom? To analyze this diversity and start to answer these questions, we cannot rely on traditional microbial phylogenetic analyses which often assume vertical inheritance and track variation with respect to a single reference genome.

Instead, we use a pangenome approach in which genomes are considered dynamic gene pools, shaped by vertical inheritance and horizontal exchange. Pangenome analyses focus on the set of all genes found in a population, and use comparative genomics to characterize the diversity and study the evolution of genomes. These analyses have highlighted hotspots of genetic transfer in the accessory genomes of various bacterial strains [15, 16, 17], as well as provided evidence for frequent recombination between phages in specific medical or environmental contexts [18, 19, 20].

Here, we reexamine the mosaicism hypothesis on a set of 3425 diverse phages, retrieved from [The Actinobacteriophage Database](https://phagesdb.org/) [21]. We aim to characterize the diversity of these genomes both at a large scale, investigating the order and structural organization of genes in the genome, and at a small scale, calculating how the linkage between mutations decays as a function of genomic distance.

### 1.1 Description of the phage database

In the Actinobacteriophage Database, the phages are organized into clusters by sequence similarity, and some are further divided into subclusters [12, 22]. Genes are sorted into gene families, known as phams [23, 24]. For our analysis, we consider subclusters and clusters that have not been subclustered, and only those which contain at least 10 phages. There are 88 such groups, as we will refer to them henceforth, which contain a total of 3467 phage genomes. Downstream analysis was restricted to 3425 phages across 86 groups; further information on the data is available in Methods 4.1.

The groups are heterogeneous with respect to size, diversity, morphotype, life cycle, and isolation host; a summary of these characteristics can be found in Supplementary Information (SI) A. Notably, the groups do not represent a single taxonomic level, but instead vary widely in diversity, as displayed in Fig. 1.

**Figure 1:**
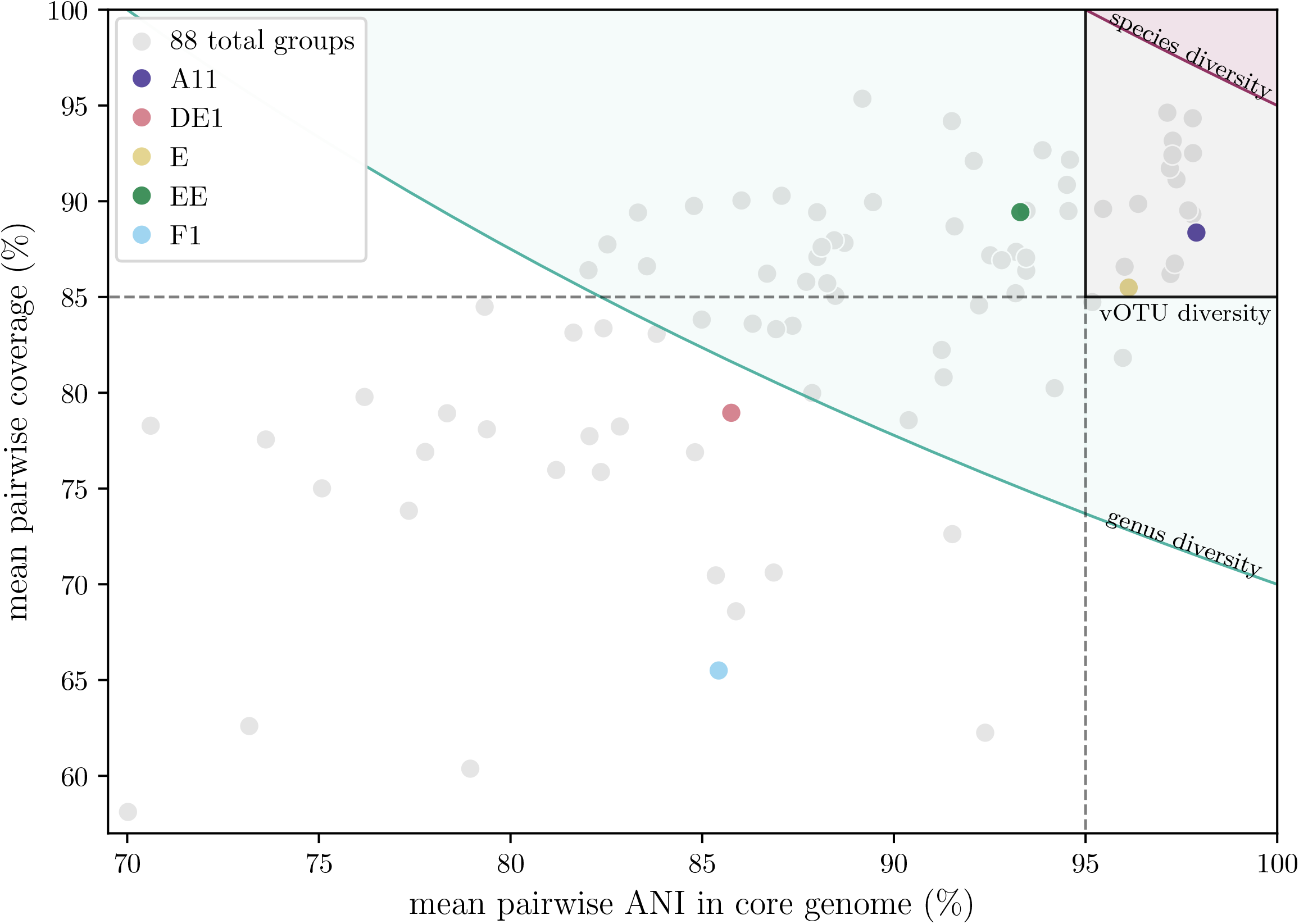
Diversity within different groups. The pairwise average nucleotide identity (ANI) is calculated from the pairwise Hamming distance in the core genome alignment, which is illustrated in Fig. 2 a). The pairwise coverage is calculated by taking the percentage of the smaller genome covered by pairwise shared phams. These measurements are a proxy for the vOTU determiners of ANI and coverage; the approximate vOTU thresholds are in black, and the groups in the gray-shaded region have diversity approximately consistent with that of vOTUs. The approximate thresholds for species and genera are the pink and teal lines respectively. The groups consistent with diversity of a genus are in the teal-shaded region. The mean pairwise ANI across the whole genome is approximated by the mean pairwise coverage multiplied by the mean pairwise ANI in the core genome. The groups that are colored are referenced in later figures.

Some groups contain closely related phages that could be considered comparable to a viral operational taxonomic unit (vOTU), a classification often used in metagenomic studies with a recommended criteria of 95% average nucleotide identity (ANI) over at least 85% of the shorter genome [25]. With respect to the recommended taxonomic criteria of species and genera, 95% and 70% nucleotide identity of the entire genome respectively [8], some groups could be considered a genus, but none would be considered a species. Group EE is an example of a genus-like group, colored in green in Fig. 1. Other groups contain much higher diversity and might be more appropriately compared to a taxonomic family, such as group F1, colored in light blue in Fig. 1.

### 2 Results

### 2.1 Synteny is strongly conserved

The pangenome of each group, the set of all of phams present in the group, can be visualized using a pangenome graph, illustrated with examples in Fig. 2. In the pangenome graph, each pham is a node, and each genome, viewed as an ordered list of phams, is represented as a path through the phams. The pangenome graph allows us to visualize many genomes simultaneously and easily distinguish between the core phams, defined as those which appear exactly once in all phages in the group, and the accessory phams.

**Figure 2:**
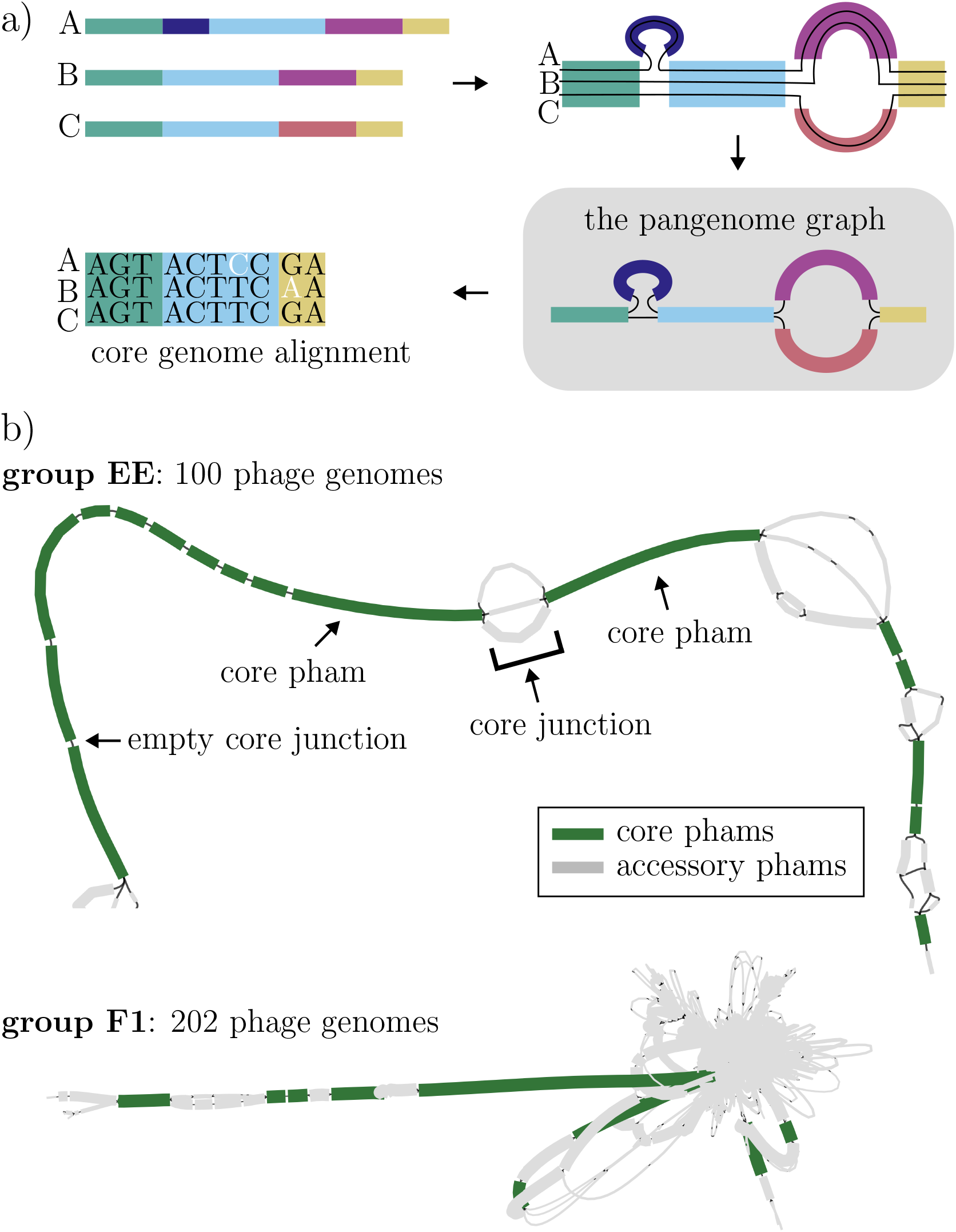
a) Illustrative diagram of a pangenome graph. Each colored block represents a different pham (gene family) and is a node in the graph, and each individual phage genome is a path through the blocks. The core genome alignment is then constructed from the alignment of all of the core phams, the phams which appear exactly once in every phage of the group. b) The pangenome graphs for group EE and F1. Group EE has 100 phages containing 44 phams, 17 of which are core. Group F1 has 202 phages containing 863 phams, 9 of which are core. Core phams are shown in green, and accessory phams in gray. The length of each pham is representative of the average length of all of the genes in the pham, and the thickness of each pham represents how many genomes contain that pham.

Pangenome diversity differs widely among groups in the database. Two examples, group EE and group F1, are shown in Fig. 2 b). The genomes in group EE have more similar pham content than those in group F1, but in both groups, the core phams appear in the same order in every phage in the group. 78 out of the 88 groups exhibit complete synteny in the ordering of the core phams, as described in Fig. 3 a). In the remaining ten groups, the majority of the phages follow the consensus ordering of core phams, and even in the phages that do not, the differences in the core pham orderings are minor, as shown in Fig. 3 b).

**Figure 3:**
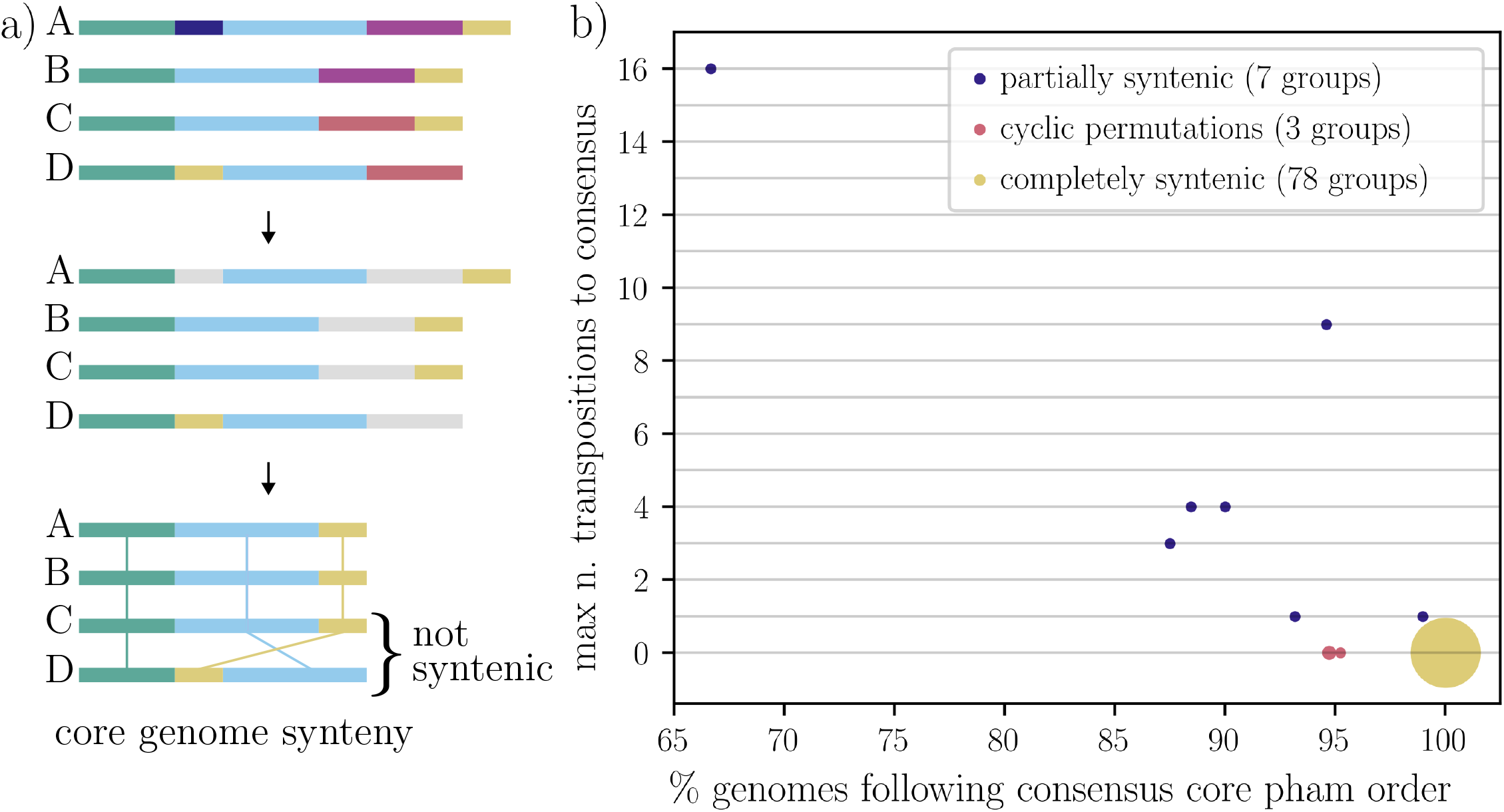
a) Illustrative diagram of synteny in the core genome. b) The extent of core synteny across the 88 groups. The *x*-axis shows the percentage of phage genomes that follow the consensus (majority) core pham order. The *y*-axis is the maximum distance between the observed non-consensus orderings and the consensus ordering for each group. For each observed non-consensus ordering, we find the minimum number of transpositions to recover the consensus ordering, and then plot the maximum across all the non-consensus orderings (usually only one, see SI B.1) for each given group. The distance metric used is equivalent to a Cayley distance. For the pink circles, the multiple core pham orderings were cyclic permutations of each other. The sizes of the circles are proportional to the number of groups represented by each circle.

Synteny extends beyond the core phams into the accessory genome. The conserved core pham order provides a natural frame of reference to define the location of accessory genes. Each accessory gene resides in a *core junction*: the genomic region between two sequential core phams, as illustrated in Fig. 2 b). We then can ask whether genes from the same pham appear in the same junction, and we find that for almost all of the accessory phams that appear in more than one phage, their genes are always found in the same core junction; the distribution is shown in SI Fig. S2 a). The order conservation also extends to within junctions; the pairwise ordering of accessory phams within the same junction is strongly conserved, with more details in SI Fig. S2 b).

### 2.2 Accessory phams are localized

To investigate the distribution of accessory phams along the core genome backbone, we exclude phages that do not conform to their group’s consensus core synteny. This restriction excludes less than 1% of the phages overall and only two entire groups (see Methods 4.1 for more details). The resulting 86 groups have perfectly syntenic core genomes, ensuring that each accessory gene resides in a single well-defined core junction.

We illustrate the distribution of accessory phams in group EE in Fig. 4 a), by plotting the number of accessory phams located in each junction. We find some junctions to be “hotspots”, containing many accessory phams, while the majority are empty. In group EE, over a third of the accessory phams are located in just one junction (#12), over half of the accessory phams are located in two junctions (#12 and #16), and all of the accessory phams are in only a third of the junctions, leaving two thirds of the junctions empty.

**Figure 4:**
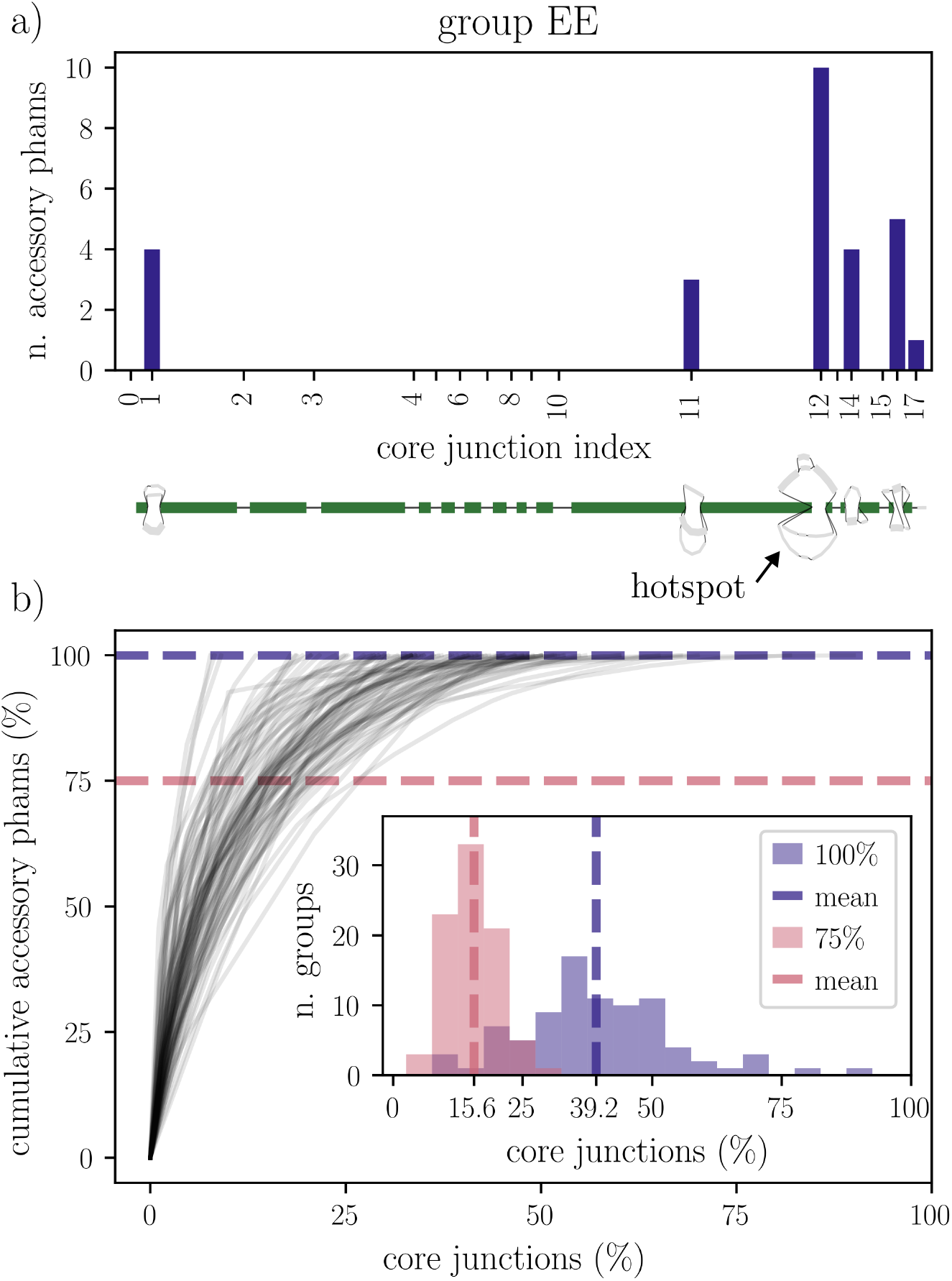
Localization of accessory phams. a) Distribution of the location of the accessory phams for group EE. The pangenome graph is shown below for reference. b) Lines indicate the (minimum) percentage of core junctions required to hold *y* percent of the accessory phams. There is one line per group. For example, the point (10, 50) would illustrate that 10% of the core junctions contain 50% of the accessory phams. The dotted horizontal lines correspond to the inset which shows the percentage of core junctions required to contain 100% (indigo) and 75% (pink) of the accessory phams respectively. The unweighted means across all of the groups are shown with the vertical dotted lines. Panel b) is inspired by [15].

We can formalize these statistics by calculating the minimum percentage of junctions required to contain a given percentage of accessory phams. In Fig. 4 b), we quantify this over all of the groups. The general pattern of EE is found in all of the groups; accessory phams are strongly localized. Most of the core junctions are empty, and most of the accessory phams are located in a few core junctions. On average across all of the groups, 75% of the accessory phams are found in only around 16% of the possible junctions.

One possible factor contributing to keeping core junctions empty is the overlap of sequential core phams. On average per group, just over half of the empty junctions coincide with overlapping flanking core phams; see SI Fig. S4 for more details. The localization of the accessory phams along the core genome can also be quantified with measures such as entropy and participation ratio. Both of these analyses provide further evidence for significant localization, which is shown in SI Fig. S3.

A naturally arising question is if there are any functional signatures of the “hotspots”, or their flanking core phams. Accessory phams involved in recombination and defense are almost always found in hotspots instead of elsewhere in the accessory genome. Also, a higher proportion of core phams flanking hotspots had anti-defense functions when compared to the proportion of all core phams. A full analysis is in SI C.3 and SI Fig. S5, and while these anecdotal results are biologically interesting, the percentage of phams with annotations is currently too low to make any sweeping statistical statements.

### 2.3 Linkage decay in the core genome provides evidence for homologous recombination

The analysis of the accessory genome and the presence of accessory “hotspots” suggest substantial genetic exchange between phages. One may therefore expect that phages also exchange homologous genetic material in their core genomes. To investigate this, we calculate linkage disequilibrium, a measure of the correlation between two single nucleotide polymorphisms (SNPs) in the core genome alignment, which is illustrated in Fig. 2 a). With frequent recombination, we expect linkage disequilibrium to decay with distance between SNPs, as the probability that the alleles came from the same recombination block would decrease with distance. With no recombination, we expect evolution to be tree-like and linkage to be independent of distance. Details of the linkage disequilibrium calculation can be found in Methods 4.4.

Linkage disequilibrium decays as a function of genomic distance in the core genome with power-law scaling to a “residual linkage” level (which differs from group to group) as shown for three different groups in Fig. 5. This observed decay of local linkage disequilibrium provides evidence of frequent and widespread homologous recombination between related phages.

**Figure 5:**
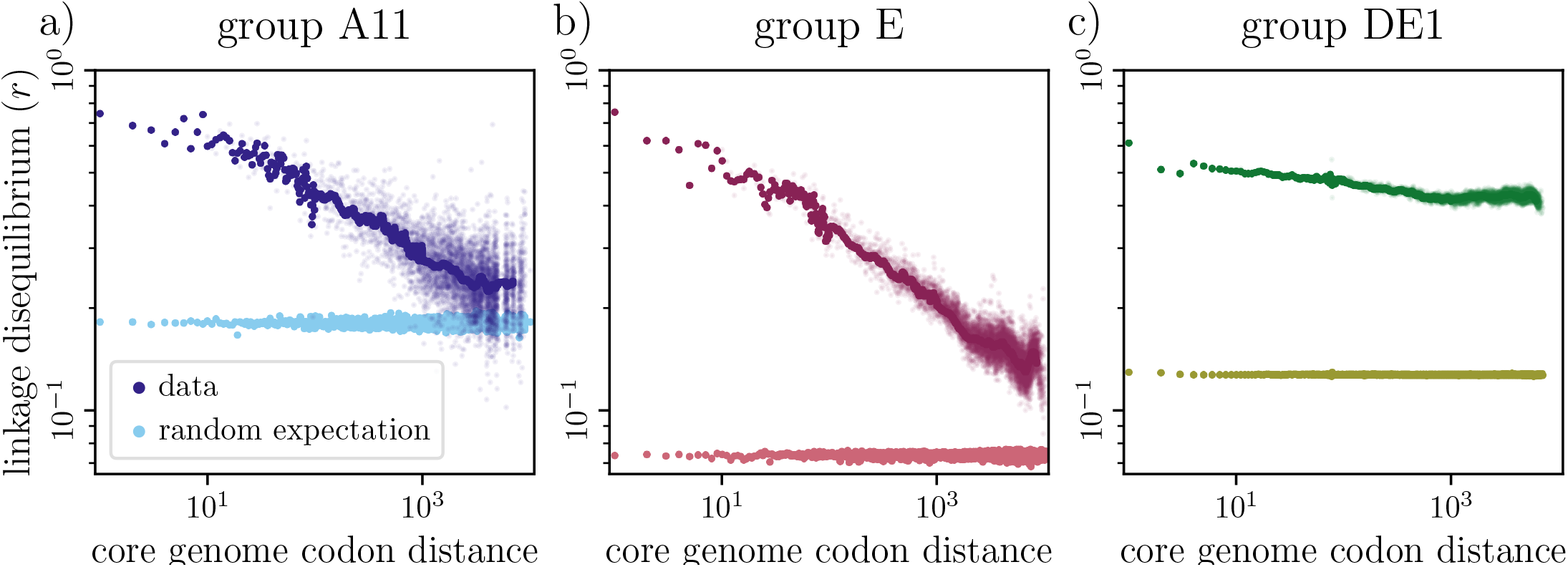
Linkage disequilibrium decay with distance along the core genome for three phage groups. Values are compared to the background linkage that would be observed even if mutations were perfectly unlinked (random expectation). This is calculated by reshuffling each column of the nucleotide alignment separately (see Methods 4.4 for more details). The linkage disequilibrium average at each distance is weighted by SNP density to ensure uniform contributions across the core genome from regions of both high and low SNP density. Further details are available in SI D.2. a) Group A11 shows rapid decay of linkage with distance in the core genome, reaching background random expectation. b) Group E shows similar behavior. c) Group DE1 decays at a slower rate and asymptotes to a residual linkage value much higher than the background random expectation. The smaller dots are all data points, and the larger dots are a rolling average.

While for around 20% (17/86) of the groups analyzed, including groups A11 and E in Fig. 5 a)-b), linkage disequilibrium at long distances decays to the level of detectability (defined quantitatively in SI E.2), the majority of the phage groups exhibit a varying degree of genome-wide residual linkage, as shown in SI Fig. S8 a). Some groups, such as group DEI illustrated in Fig. 5 c), show high residual linkage across the entire core genome, pointing to the existence of phylogenetic structures within these groups of genomes, that persist despite ongoing recombination.

The linkage decay indicates that recombination is acting on short length scales, so we quantified relevant distances with further genomic analyses. Measuring the distance between phylogenetically incompatible SNPs, a signature of recombination, we found the mean distance per group to be on average 66 base pairs (see SI F for more details). This is smaller than would be expected by recurrent mutations, as shown in SI Fig. S9. To estimate an upper bound on the recombination block size, we considered single recombination events evident by regions of high divergence in pairs of closely related phages, as in SI Fig. S10. The mean of the maximum length of these regions per group is 570 base pairs, which is on the order of the length of a core pham, as shown in SI Fig. S11.

### 2.4 Long-range linkage reveals phylogenetic substructure

The groups with higher residual long-range linkage tend to exhibit more pronounced phylogenetic substructure. Considering all pairwise Hamming distances in the core genome alignment, hierarchically clustering the phages reveals a nested block structure in the distance matrices of all the groups, as shown in the insets in Fig. 6. Across groups, as the amount of residual long-range linkage varies, so does the extent which the nested blocks have diverged. Groups for which the matrix can be partitioned into a clear block structure are the ones in which it is possible to meaningfully subdivide groups into well-defined subgroups. Such groups exhibit subgroup-specific haplotypes extending the length of the core genome, which creates the long-range linkage disequilibrium.

**Figure 6:**
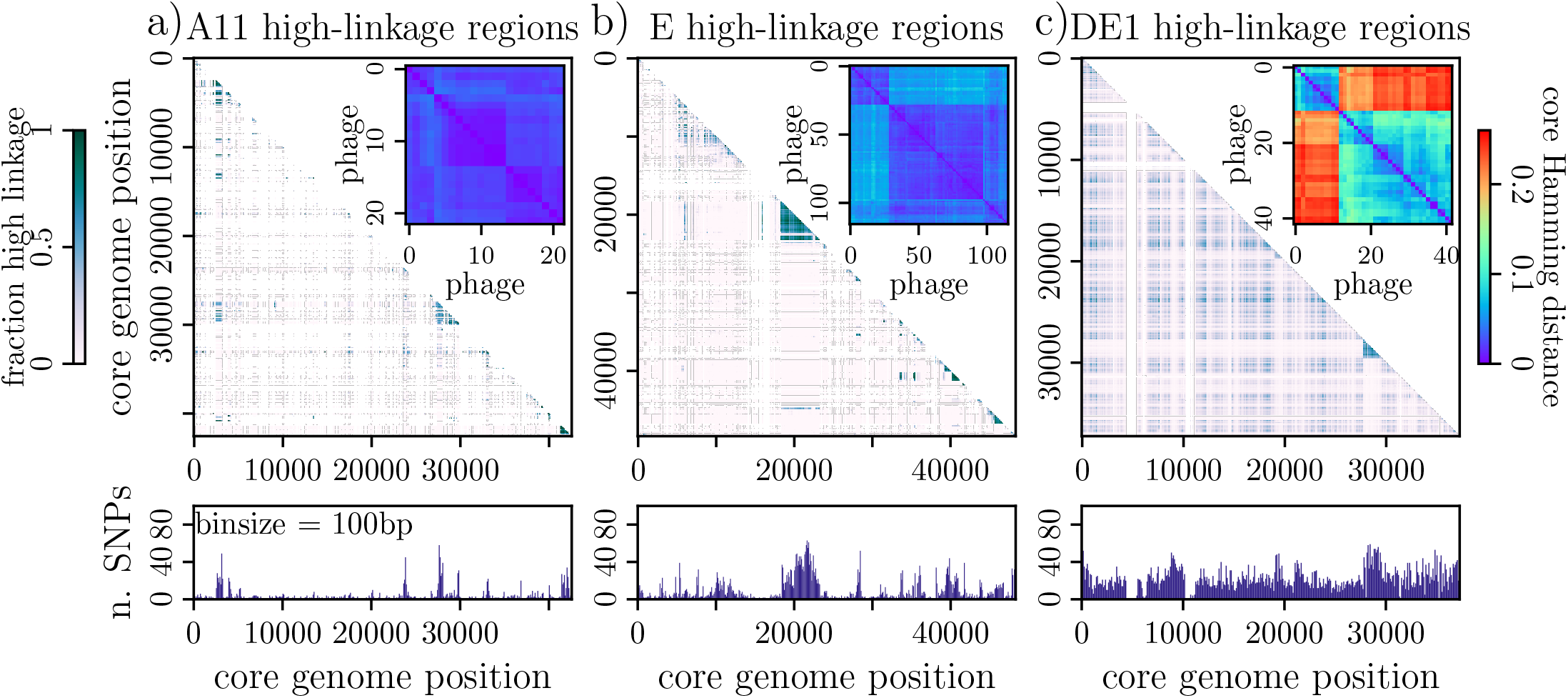
a)-c) upper show the linkage enrichment plots, where bin sizes of 100bp by 100bp are colored according to the proportion of pairs with linkage in the top 90th percentile. White bins indicate no data, due to no SNPs. a)-c) lower show the SNP density across the core genome with a bin size of 100bp. a)-c) inset show the distance matrix for the pairwise core Hamming distances for each pair of phages in the group. a) The phages in group A11 are closely related, with few SNPs across the core genome. High linkage is predominantly near the diagonal of the linkage enrichment plot. b) In group E, there is one region in the core genome that has high linkage and a high SNP density. This is caused by the region separating into two distinct haplotypes. This also causes a slight block structure in the distance matrix. c) Group DE1 shows a distinct subgroup structure in the distance matrix. This results in high SNP density and linkage across the entire core genome.

An example is group DE1, as illustrated in Fig. 6 c); SNP density is high across most of the core genome (bottom), there is elevated linkage disequilibrium across the entire genome (top), and pairwise Hamming distance calculations reveal two distinct subgroups (inset). If we consider the two haplotypes separately and analyze the linkage disequilibrium of the two subgroups individually, we recover steep, rapid decay that roughly reaches the random unlinked expectation at long distances, as shown in Fig. 7. This indicates that recombination is more frequent within each of the subgroups.

**Figure 7:**
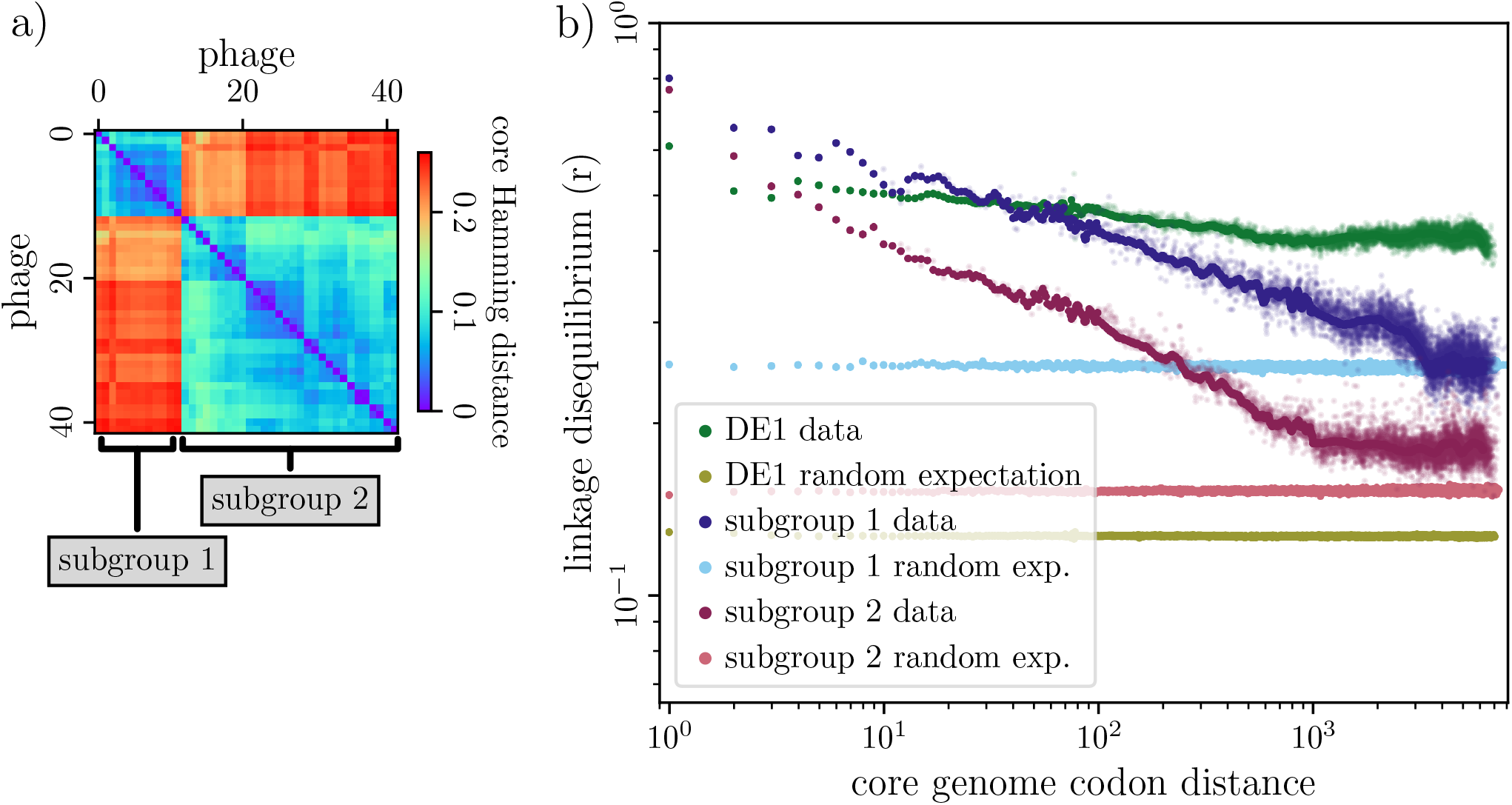
a) The distance matrix for the pairwise core Hamming distances for each pair of phages in group DE1, with brackets denoting the two subgroups. b) The linkage disequilibrium of group DE1 (greens) and its two subgroups separately (blues and pinks).

In some groups, we find this phylogenetic structure in only one small region of the core genome. This is shown in Fig. 6 b) for group E; a small region around core genome position 20000 has a high SNP density and has high linkage disequilibrium within that region. In this region, the core genome alignment has split into multiple haplotypes. We identified this phenomenon in 13 of the 86 groups analyzed. For these groups, we also analyzed the haplotypes separately by only considering SNPs that were deviations from the haplotype consensus sequence in that region. This also recovered steeper linkage decay that reaches closer to the unlinked random expectation, shown in SI Fig. S7 for group E.

These results show that the patterns of linkage disequilibrium are heavily influenced by phylogenetic structure and spatial heterogeneity across the core genome. The spatial heterogeneity may be indicative of a region containing a recent recombination event, or alternatively a region that has sufficiently diverged such that homologous recombination is no longer possible. Moreover, these results indicate that recombination is rampant within the relevant phylogenetic substructures.

## 3 Discussion

In this paper, we analyzed a large dataset of diverse actinobacteriophages which had been clustered into groups of varying sizes and diversity. We characterized these groups and considered each of their pangenomes to investigate genome structure and linkage disequilibrium within each group. We found strong synteny in the core and accessory genome, with accessory phams localized into hotspots. In some groups, we found evidence for extensive homologous recombination in the core genome across the entire group, while in others we found a clear separation into subgroups, with recombination more prevalent within subgroups than across.

These findings of conservation and restriction in addition to frequent recombination are two sides to the same coin. The rapid decay of linkage disequilibrium indicates frequent shuffling of genetic material. Yet, despite ongoing homologous recombination, phage groups exhibit persistent phylogenetic structure manifested by the residual long-range linkage. This could be explained either by the phage “population structure”, for example different host range, that prevents genetic exchange between certain subgroups, or by the effect of selection that amplifies certain core genome haplotypes.

The core synteny and localization of accessory phams are heavily influenced by overlapping genes. Phage genomes are strictly constrained in length due to the physical size of their capsid, so genes sharing coding sequences in phages is common and well-documented [26]. On average, half of the empty junctions in a group correspond to overlapping core genes, as shown in SI Fig. S4.

The localization of accessory phams into hotspots also agrees with the genomic organization of the phage’s bacterial counterparts, which themselves have hotspots. In bacteria, these hotspots often have specific functional traits, for example, defense islands [27]. The hotspots in phages could also potentially represent corresponding functional units such as anti-defense islands or defense-islands in temperate phages, an idea which is supported by our preliminary analysis in SI C.3. As functional annotation tools continue to improve, further work should be done to try to classify and understand the functions of not only the hotspots, but also the conserved core genes.

It is worth noting that the presented analysis relies on the pham classifications, which are not robust to database additions. Our designations of phams as “core” or “accessory” were not based on their known biological function, but rather derived from their pangenome distribution in our genomic analysis. Our findings of synteny and linkage decay are robust to inevitable slight variations in pham classifications as more phages are added to the database.

This study was focused solely on the analyses within groups. More can be learned about phage mosaicism from this dataset through investigating inter-group statistics of the phams that exist across many diverse groups. This work also opens up opportunities to use similar methods to analyze other groups of phages and takes us one step further in in our understanding of phage evolution and mosaicism.

## 4 Methods

### 4.1 Data

The data were obtained using the API of phagesDB.org [21], as well as accessing NCBI GenBank [28] via ncbi-acc-download (https://github.com/kblin/ncbi-acc-download). Data can be downloaded directly from the API and GenBank using the pipeline we created to download, filter, and sort the data: https://github.com/jfendley/phage-download. The pipeline was run on 5 March 2024 for the data shown in the paper. Only genomes from the database that had also been published in GenBank were used for the analysis.

The genes in the database had been “phamerated”: sorted into gene families, “phams” [23], with a tool, PhaMMseqs, that uses MMseqs2 [24, 29]. The pham classifications are a function of the data used and are not invariant to database additions. If one were to repeat the analyses on freshly downloaded data, the quantitative results may change slightly, but the qualitative results will be robust.

The genomes in the database have been sorted into clusters by sequence similarity. Some clusters were further divided into subclusters. A description of the initial clustering methodology can be found in [12], and recently, an automatic clustering method was shown to support the original cluster classifications [22]. For this analysis, we considered only irreducible groups of phages, meaning either subclusters, or clusters that had not been divided further into subclusters. We refer to these as “groups”.

We restricted the analysis to only groups which contained at least 10 phages. In total, 88 groups containing 3467 genomes were analyzed initially, and then phages that did not follow the consensus core gene order were removed so we could analyze further 86 perfectly syntenic groups with 3425 phage genomes. The two entire groups that we did not analyze further were the outlier in Fig. 3 b), and a group that upon removal of the non-syntenic phages no longer reached the 10 phage minimum.

These groups vary significantly in size, diversity, and biology. A brief overview of properties of the groups is available in Fig. S1, and a summary table is included in the Supplementary Material. All of the phages were actinobacteriophages and were isolated on 35 different host species (7 genera).

### 4.2 Reproducibility

All of the code for the analysis is available in the GitHub repository: https://github.com/jfendley/phage-pangenomes. Instructions and a Bash script are provided to recreate all of the analyses.

### 4.3 Core genome construction

Core phams are those that appear exactly once in every phage in the group. The core pham amino acid sequences were aligned separately using MAFFT [30]. The nucleotide sequence alignment was inferred from the amino acid alignment, with the reverse complement taken for phams encoded on the reverse strand. The core genome alignment was created by concatenating the nucleic acid alignments for each pham in the syntenic order. Core genome alignments were only created for the syntenic groups.

### 4.4 Linkage disequilibrium analysis

Considering two different loci in the core genome (1 and 2), and one specific allele per locus (A and B), linkage disequilibrium is defined as:

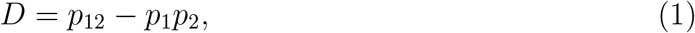

where *p*_1_ is the frequency of allele A at locus 1, *p*_2_ is the frequency of allele B at locus 2, and *p*_12_ is the frequency at which A appears at locus 1 while allele B appears at locus 2. As *D* strongly depends on the total number of phages, we instead use the correlation coefficient, *r*, defined by:

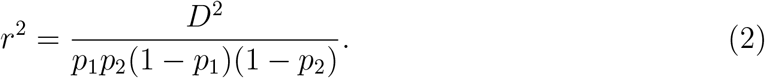

The coefficient *r* takes values between 0 and 1, but the allowed range depends upon the frequencies *p*_1_ and *p*_2_, which depend on the number of phages. For randomly shuffled unlinked alleles, the expectation of *D* is 0, but the expectation of *r* is dependent on the frequencies and hence number of phages. We assume there are *N* phages, so that *p*_1_ = *N*_1_*/N, p*_2_ = *N*_2_*/N*, and *p*_12_ = *N*_12_*/N*. The unlinked random case would have the random variable *N*_12_ follow the hypergeometric distribution with parameters *N, N*_1_, and *N*_2_. This would give *r* the following expectation:

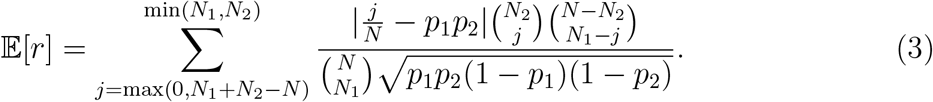

For each pair of sites, we only calculated one value of *r*, using the frequencies of the most dominant allele. For bi-allelic sites, this is comprehensive and complete, as a result of reciprocal identities of *r*. A justification into this choice is available in SI D.

We considered only sites where gaps were not the most or second most frequent allele. Given *N* phages, we also considered only sites where the most dominant allele occurred in more than one phage, and in no more than *N −* 2 phages. This was to ensure the results were not heavily influenced by SNPs that only occur in one phage.

To avoid introducing artificial linkage due to repetitive calculations, we did not analyze overlapping sites, which are sites in the core genome that corresponded to multiple phams in at least one phage. These overlapping sites could be a result of a shifted reading frame or different strandedness, and are also considered further in SI C.2.

Since we are interested in linkage disequilibrium as a function of distance, we averaged over many equidistant pairs of sites. To prevent regions with high-SNP density contributing disproportionately to the average, we weighted the *r* values by SNP density. As SNP density can be very heterogeneous across the core genome, the weighting is important to ensure the measure is representative of the entire core genome. Specific details of the weighting are available in SI D.2.

## Acknowledgments

The authors are grateful for stimulating discussions with Beatriz Beamud, Aude Bernheim, Ivana Cvijović, Marian Dominguez-Mirazo, Benjamin Good, Zhiru Liu, Aditya Mahadevan, Marcie Marston, and Eduardo Rocha. This research was supported in part by grant NSF PHY-2309135, the Gordon and Betty Moore Foundation Grant No. 2919.02, the Chan Zuckerberg Initiative DAF grant to the Kavli Institute for Theoretical Physics (KITP), and by the Swiss National Science Foundation through Grant No. 310030 188547 (to RAN).

## Supplementary Information

### A Group summary

A table summarizing key properties of all 88 groups analyzed is available in the Supplementary Material file: supplementary table.tsv. Table S1 describes each of the columns in the file. The distributions of a selection of properties are shown in Fig. S1.

**Table S1:** Description of the column names in the Supplementary Material table summarizing key properties of all 88 groups.

| Column name | Description |
| --- | --- |
| name | name of the phage group |
| type | “cluster” or “subcluster” in the database |
| cluster | which database cluster it is in (itself if a cluster) |
| life_cycle | Temperate or Lytic |
| n_phages | number of phages |
| n_phams | number of phams in pangenome |
| n_core | number of core phams |
| synteny | “syntenic”, “nonsyntenic”, or “cyclic” |
| mean_genome_length | mean length of genomes |
| mean_n_phams | mean number of phams per phage |
| mean_percent_core | mean percent of genomes by length that is core phams |
| mean_percent_genes | mean percent of genomes by length that is coding regions |
| mean_percent_pairwise_coverage | mean pairwise coverage: percent length of the smaller genome that is pairwise shared phams |
| mean_hamming | mean pairwise core genome Hamming distance |
| mean_jaccard | mean pairwise Jaccard distance of shared phams |
| std_hamming | standard deviation of pairwise core genome Hamming distance |
| std_jaccard | standard deviation of pairwise Jaccard distance of shared phams |
| most_common_morphotype | the most common morphotype |
| all_morphotypes | list of all morphotypes |
| most_common_host_genus | most common isolation host genus |
| all_host_genera | list of all isolation host genera |
| most_common_host_species | most common isolation host species |
| all_host_species | list of all isolation host species |

**Figure S1:**
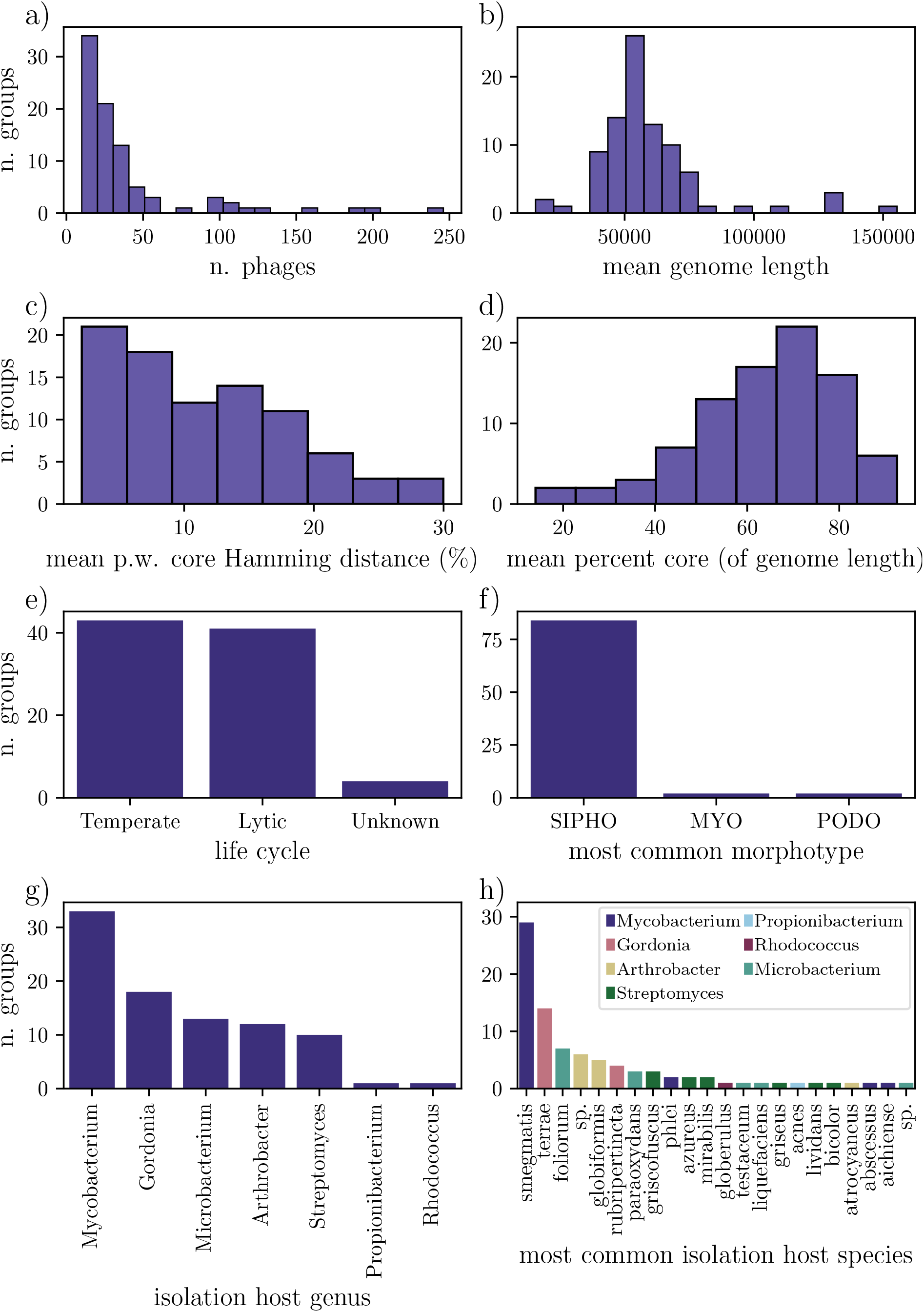
The distribution of a) the number of phages in each group, b) the number of core phams for each group, c) the mean pairwise core Hamming distance for each group, d) the mean percentage genome length of core phams for each group, e) life cycles per group, f) the most common morphotypes per group, g) the isolation host genera (exactly one per group), and h) the most common isolation species per group, colored by isolation host genera.

### B Synteny

#### B.1 Core synteny

The definition of a core pham requires the pham to appear exactly once in every phage. Some of the accessory analyses also focus on accessory phams that appear at most once in every phage. These restrictions do not exclude many phams, as less than 2% of all of the phams in the dataset appear more than once in a single phage. 40 groups have no duplicate phams, and group BE2 has the highest percentage of duplicates, with just under 14% of its phams appearing more than once in at least one phage. These numbers are sufficiently low for our analytical purposes.

**Table S2:** The groups with non-syntenic core genomes and a description of their pham order differences. The “N. orderings” column describes the number of core genome orderings present in all of the phages. The “Phage split” column details the number of phages in the most frequent ordering, and then the number of phages in the subsequent orderings. The “Description of differences” column details how the multiple orderings are related to each other. “Flips” means the strandedness of the pham changes.

| Group | N. phages | N. orderings | Phage split | Description of differences |
| --- | --- | --- | --- | --- |
| AO2 | 16 | 2 | 14, 2 | one pham moves two positions |
| AR | 19 | 2 | 18,1 | cyclic permutation |
| B3 | 44 | 2 | 41,3 | one pair of phams switch places |
| BG | 10 | 2 | 9, 1 | one pair of phams switch,<br>of which one flips |
| CA | 38 | 2 | 36,2 | cyclic permutation |
| CR2 | 21 | 2 | 20,1 | cyclic permutation |
| EC | 37 | 3 | 35,1,1 | one pham travels to<br>two different locations |
| EF | 26 | 2 | 23,2 | one pham moves three positions |
| K1 | 99 | 2 | 98,1 | one pair of phams switch places |
| K6 | 21 | 2 | 14,7 | one pham moves 16 positions |

A description of the groups with imperfect core synteny is available in Table S2. All of the GenBank files describe AR, CA, and CR2 as linear. However, for AR, the information from the Actinobacteriophage Database described the end types as “circular”. The cyclic permutations could have been a result of assembly errors, and circularization of DNA has been shown during a phage life cycle [31].

We continued our analysis by cyclically permuting the order of the phams in the outlier phages in groups AR, CA, and CR2, and neglecting the outlier phages in AO2, B3, EC, EF, and K1. We chose to not analyze BG or K6 any further, since if we neglected the outlier in BG it would no longer meet our threshold size of at least ten phages per group, and the outliers were too great a proportion of the phages in K6.

#### B.2 Accessory synteny

We set out to determine if accessory phams always appear in the same core junction if they appear in different phage genomes. We exclude the phams that are duplicated in at least one phage, see SI B.1. In over half (47/86) of the groups, all of the accessory phams always appear in the same core junction. In all other groups, a high percentage (at least 93%) of the accessory phams appear in exactly one core junction, see Figure S2 a). Accessory phams are hence individually localized; when they appear, they usually appear in the same context within the core backbone.

To analyze the synteny within the accessory genome, we looked at the synteny within each non-empty junction. We took all pairs of phams in that junction that appear together in at least two phages, and ask if in these cases they always appear in the same order. In 43% (37/86) of the groups, all of the junctions were perfectly syntenic, as shown in Fig. S2 b) *x*-axis.

In the 49 groups that do not have perfect accessory synteny, less than half of the junctions were not-syntenic, see Fig. S2 b) *x*-axis. Within those non-syntenic junctions, typically less than a third of the co-occurring accessory pham pairs were not syntenic (appeared in multiple orderings), as shown in Fig. S2 b) *y*-axis. These results suggest that there is strong order conservation within the accessory genome.

**Figure S2:**
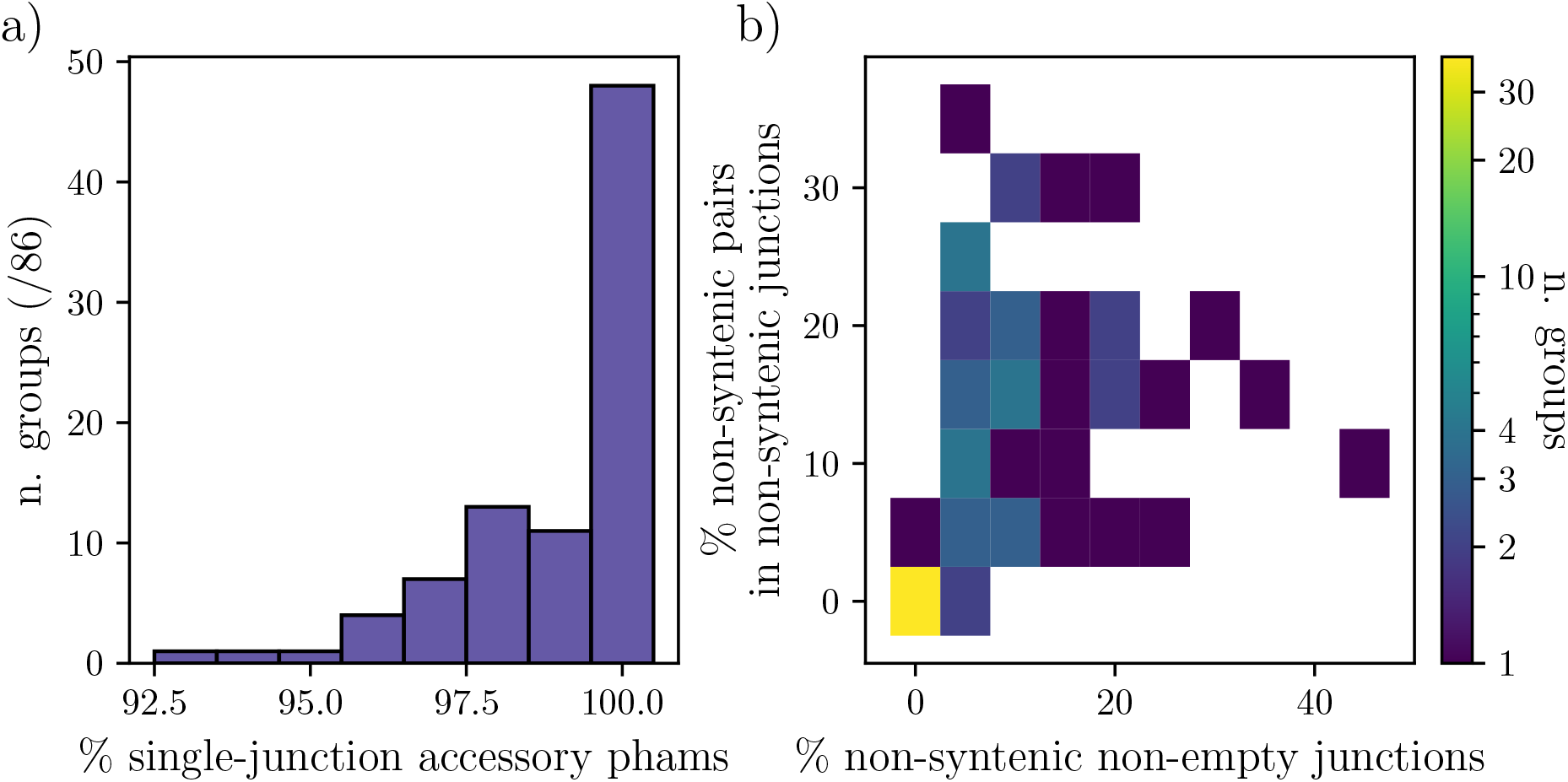
Accessory genome synteny. a) The distribution across groups of the percent of multi-phage accessory phams that always appear in the same junction. In 47 groups, all of the accessory phams always appear in the same core junction. b) Pairwise accessory synteny in core junctions. Considering non-empty junctions, we compared the pairwise ordering of accessory phams. The *x*-axis shows the distribution across groups of the percent of non-empty junctions which do not have pairwise accessory synteny. 37 groups have perfect pairwise accessory synteny. For the remaining 49 groups, the *y*-axis shows the distribution of the percent of all pairs of accessory phams that are non-syntenic.

### C Localization of accessory phams

#### C.1 Measures of localization

Each of the *N* total core junctions, labeled *i*, contains a (potentially empty) fraction *p*_*i*_ of all of the accessory phams. The entropy is thus calculated as

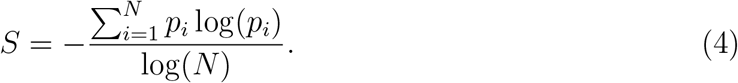

The participation ratio is defined as:

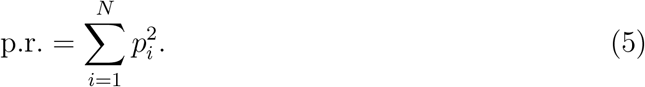

This has the property that if the accessory phams are perfectly equally distributed then p.r. = 1*/N*. Moreover, if they are equally distributed in *n* core junctions, where *n* ≤ *N* core junctions, then p.r. = 1*/n*.

The normalized participation ratio is defined as:

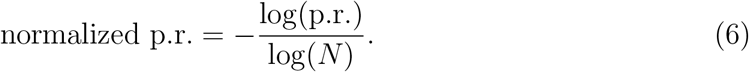

All three statistics are shown in Fig. S3 and indicate significant localization.

**Figure S3:**
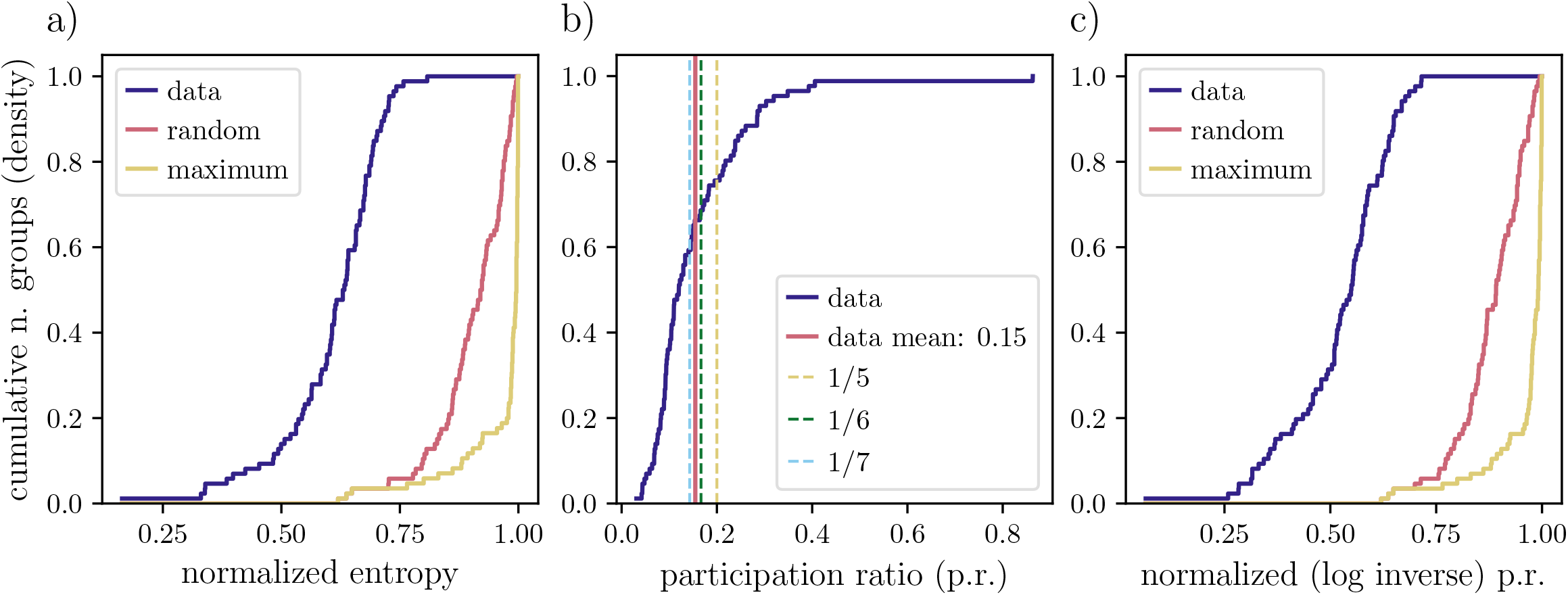
Measures of localization of accessory phams along the core genome backbone. a) The indigo line shows the density of the cumulative distribution of the normalized entropy for each of the groups. The pink line shows the distribution if each accessory pham were placed randomly in a core junction with equal probability. The yellow line shows the distribution of the maximum possible entropy if all of the phams were as evenly distributed as possible. The difference between these distributions shows that the accessory phams are significantly localized along the genome. b) The indigo curve is the density of the cumulative distribution of the participation ratio, and the vertical pink line is the mean of the distribution. The mean lies between 1/7 and 1/6, which indicates that the participation ratio is roughly equivalent to if all of the accessory phams were distributed equally among only 6 or 7 core junctions on average per group. c) The normalized participation ratio shows similar results to the normalized entropy.

#### C.2 Overlapping core genes

We also investigated the effect of overlapping core genes on both synteny and the localization of accessory phams. On average, half of the empty junctions have overlap in the flanking core phams in at least one phage. This helps to explain some of the accessory pham localization, but by itself is not sufficient to explain all of the empty junctions.

**Figure S4:**
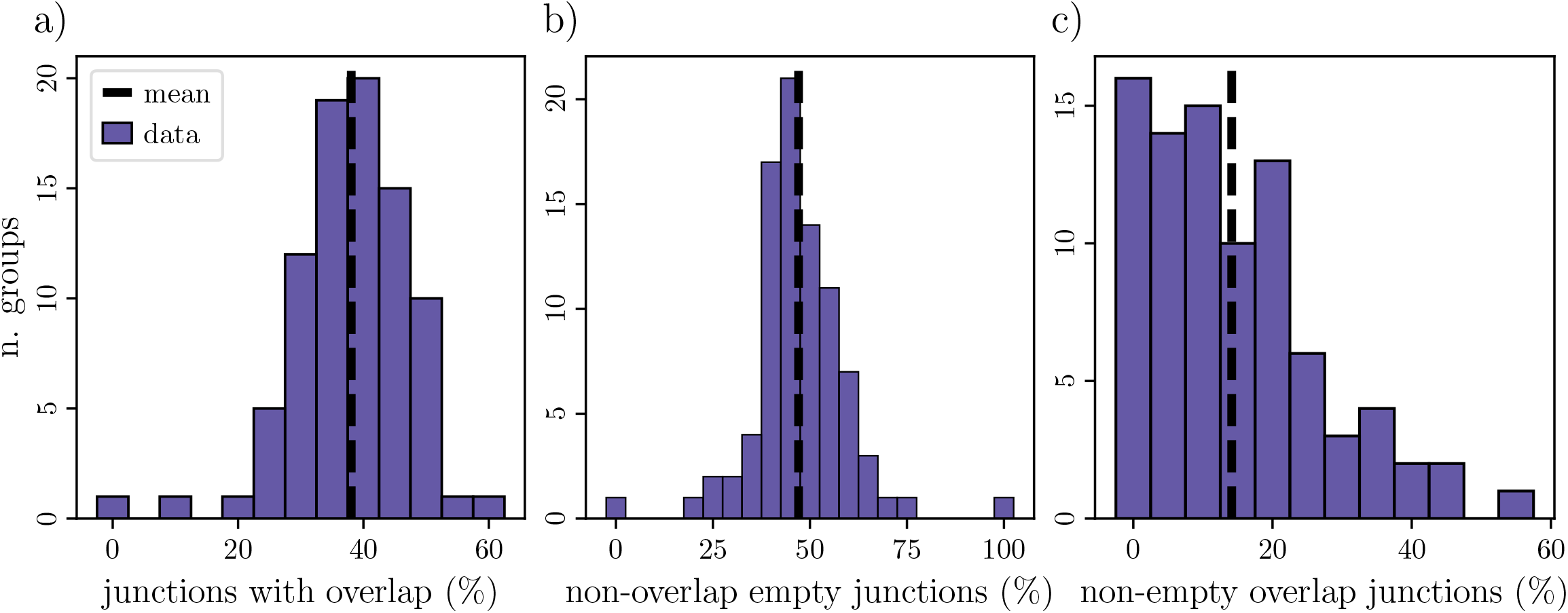
Analysis of overlapping core genes. Panel a) shows the percentage of junctions that contain an overlap (shared DNA sequence) between the flanking core phams in at least one phage. Panel b) shows the percentage of empty junctions (no accessory phams) that do not have any overlap between the flanking core phams in any phages. On average per group, just under 50% of the empty junctions do not have any phages with overlapping flanking core genes, which means just over 50% of the empty junctions do have at least one phage with overlapping flanking core genes. Panel c) shows the percentage of overlapping core phams that are flanking a non-empty junction. In other words, in at least one phage they overlap, and in at least one other phage they have an accessory pham in between them.

#### C.3 Hotspot pham functions

We are interested in patterns in the functions of genes near and in highly-occupied junctions, or ‘hotspots’. We define a ‘hotspot’ to be a junction with at least five accessory phams, chosen so that each group has at least one hotspot. Considering the (at most) four hotspots per group with the highest number of accessory phams, we look at the functions of the accessory phams within the hotspot and of the core phams flanking either side.

The database included annotations, but only 17% of phams were annotated. Here, we consider a pham to not be annotated if in all groups the pham is present in, the most common “annotation” across the genes in the pham is no annotation. Inspired by defense-islands in bacteria [27], we were particularly interested in looking for defense and anti-defense genes, so in addition to the database annotations, we used DefenseFinder [32, 33] to locate defense and anti-defense genes.

The results are in Fig. S5, comparing the enrichment of functions of core phams flanking hotspots as compared to all core phams, and likewise the enrichment of functions of accessory phams in hotspots as compared to all accessory phams. The enrichment of a function in/flanking hotspots is defined by considering the proportion of all accessory/core phams with that function (expected) to the proportion of accessory/core phams in/flanking hotspots with that function (observed). The statistic is then calculated as (observed - expected)/expected.

There is strong enrichment of integrases and G-I-Y I-Y-G endonucleases within hotspots, consistent with the hypothesis that hotspots are hotbeds for recombination [34]. There is also strong enrichment of defense genes within hotspots and slight enrichment of anti-defense genes in the core genes flanking hotspots, potentially supporting the idea of defense/anti-defense islands in phage genomes. Further investigation of hotspot functions remains an exciting avenue for future work.

### D Linkage disequilibrium

#### D.1 Bi-allelic assumption

When calculating linkage disequilibrium, we considered for each site in the core genome one dominant allele and one minority allele. This calculation is exhaustive when the site is bi-allelic. If the site is not bi-allelic, further linkage disequilibrium quantities could be calculated for the different minority alleles at the same site.

Since we are primarily interested in the effect of distance on linkage disequilibrium, it was sufficient for our analysis to only consider one linkage disequilibrium statistic per site. Moreover, the majority of all sites analyzed (65%) were bi-allelic with no gaps, and many of the multi-allelic (greater than two alleles) sites arose from phylogenetic substructure. To illustrate, groups with haplotype structure are more likely to be multi-allelic, especially if there are more than two haplotypes. When we process the haplotypes as separate subgroups (see Fig. 7), the proportion of multi-allelic sites indubitably decreases.

#### D.2 Weighting and distances

We were interested in a measure of linkage disequilibrium across the entire core genome, not skewed by regions of high SNP density. To ensure equal contributions across the core genome, we weighted each of the positions by SNP density. The weight is inversely proportional to the density of SNPs in a sliding 399 bp window, with the beginning and end of the core genome as edge cases.

Given the length of the core genome alignment, *n*, each position *I* ∈ [0, *n* − 1] was assigned a weight,

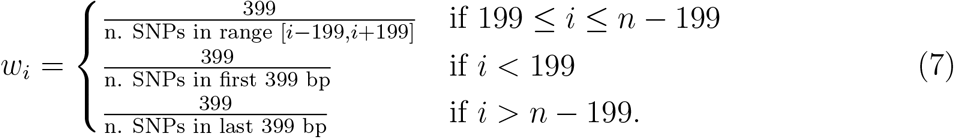

Additional window sizes were tested, but the key results remained. The window size of 399 was specifically chosen as a multiple of three to ensure equal codon-position representation in the window.

Each linkage statistic has a weight which is the product of the weights of each of the two positions considered. When looking at linkage as a function of distance, a weighted average was taken across all of the statistics with the given distance. The effect of the weighting is shown in Fig. S6. We use codon distance instead of base pair distance to average out biases due to codon position. For example, third codon positions are usually more diverse, since mutations there are more likely to be synonymous.

In the main text figures, we only consider distance within the core genome, which consists of the core phams joined together. This may not reflect the actual genomic distance between SNPs, as there may be accessory genes in between the core phams. To ensure that our results are robust, we can project the results onto a reference phage to calculate an actual distance between the two sites. Since the linkage decays so quickly on a scale comparable to the length of a core pham, projecting onto a reference genome does not change any of our main results, as evident in Fig. S6.

**Figure S5:**
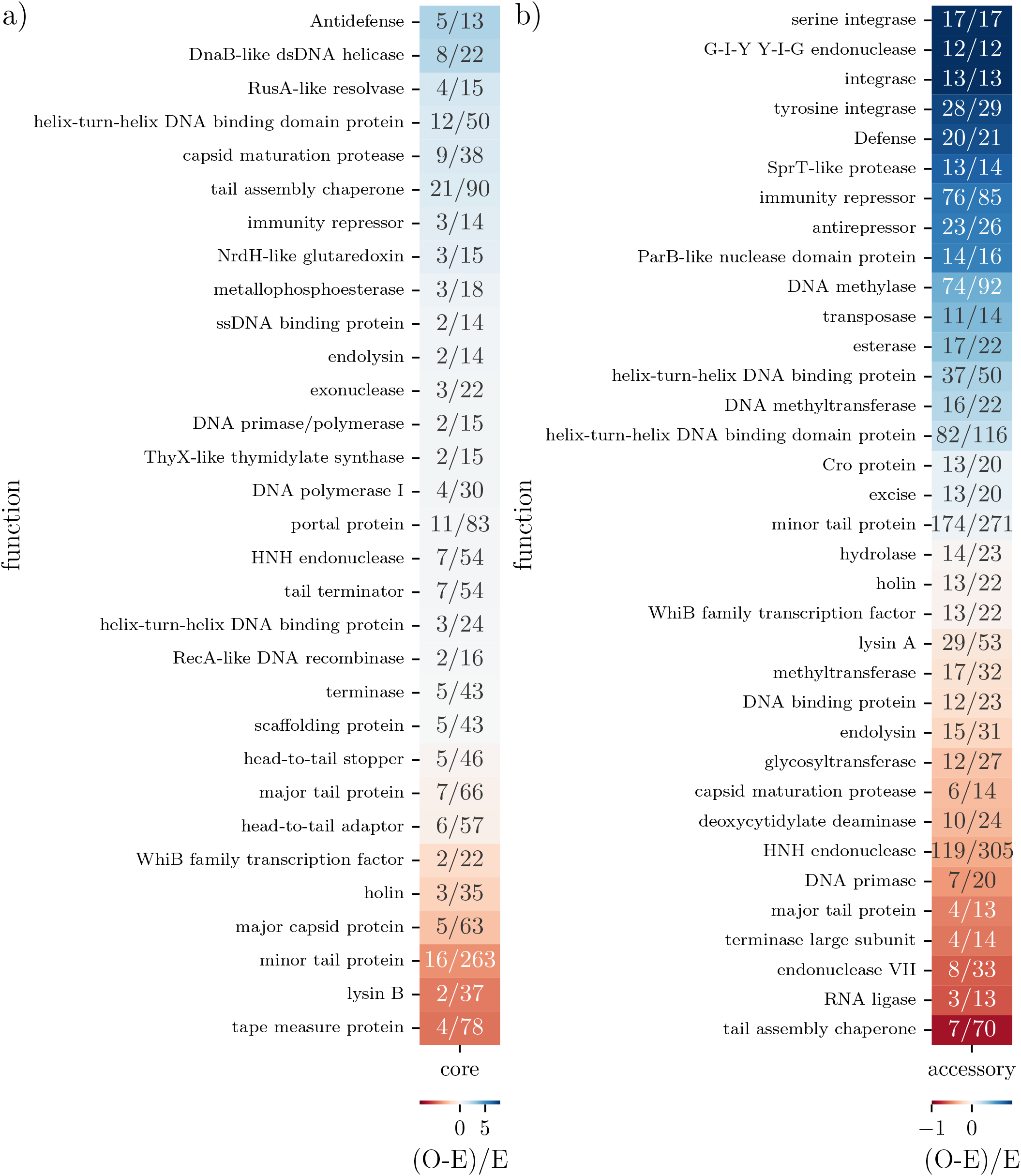
a) Heat map showing enrichment of functions of core phams that flank hotspots (observed: O) as compared to all core phams (expected: E). The numbers in the boxes are the number of core phams that flank hotspots with that function / the number of core phams with that function. b) Heat map showing enrichment of functions of accessory phams in hotspots (observed: O) as compared to all accessory phams (expected: E). The numbers in the boxes are the number of accessory phams in hotspots with that function / the number of accessory phams with that function. The scales in the color bars are chosen to reflect the maximum and minimum possible enrichment statistics, with zero in the center of the color scales.

### E Phylogenetic substructure

#### E.1 Multiple-haplotype region

We can process the groups that have regions with high linkage due to multiple haplotypes, by only considering SNPs that are deviations from the haplotypes in that region. The result is a smoothing and steepening of the linkage disequilibrium curve, as well as a lower asymptotic value, closer to the unlinked expectation. It is worth noting that the effect of the high linkage multiple-haplotype region on the original linkage disequilibrium curve is already muted since that region has high SNP density and thus has a lower weight (see SI D.2), so the processing does not cause major changes, as shown in Fig. S7 for group E.

#### E.2 Classification of groups

In around 20% of the groups (17/86), the linkage decays roughly to the random expectation. To define that rigorously, we quantified the amount of residual linkage disequilibrium at long-distances compared to the random unlinked expectation. We considered both the absolute difference between the residual linkage asymptote and the random unlinked expectation, referred to as “residual linkage (difference)”, as well as the percentage difference, referred to as “residual linkage (% difference)”.

The percentage difference was calculated as the distance between the asymptote to the average random unlinked expectation divided by the distance from the maximum weighted linkage (within codon distance ten) to the average random unlinked expectation. These statistics are shown in Fig. S8 a), and the 17 aforementioned groups are those that have a residual linkage less than 0.1 and a residual linkage percentage less than 20%.

By manual inspection of the spatial distribution of linkage disequilibrium along the core genome, as in Fig. 6, we identified 13 groups that have small high-linkage haplotype regions in the core genome. Some of these groups are included in the 17 groups that decay fully, some are not. We refer to these groups as groups with a ‘small haplotype region’.

We also manually looked at the distance matrices, as in Fig. 7 a), to identify groups that have distinct subgroup structure. 31 groups, including all 17 groups with linkage disequilibrium that decays fully and some of the small haplotype region groups, did not have (by eye) distinct subgroup structure. We refer to these groups as groups that have ‘not distinct subgroups’.

We investigated quantitative characteristics of the groups that might help determine these manual classifications. There is some correlation with the classifications and the residual linkage calculations, as shown in Fig. S8 a), as well as statistics of the distance matrix. Specifically, there appears to be a correlation with the distinctness of subgroups and the standard deviation of core Hamming distances, as well as the standard deviation of the branch lengths of the dendrogram of the distance matrix, as shown in Fig. S8 b).

**Figure S6:**
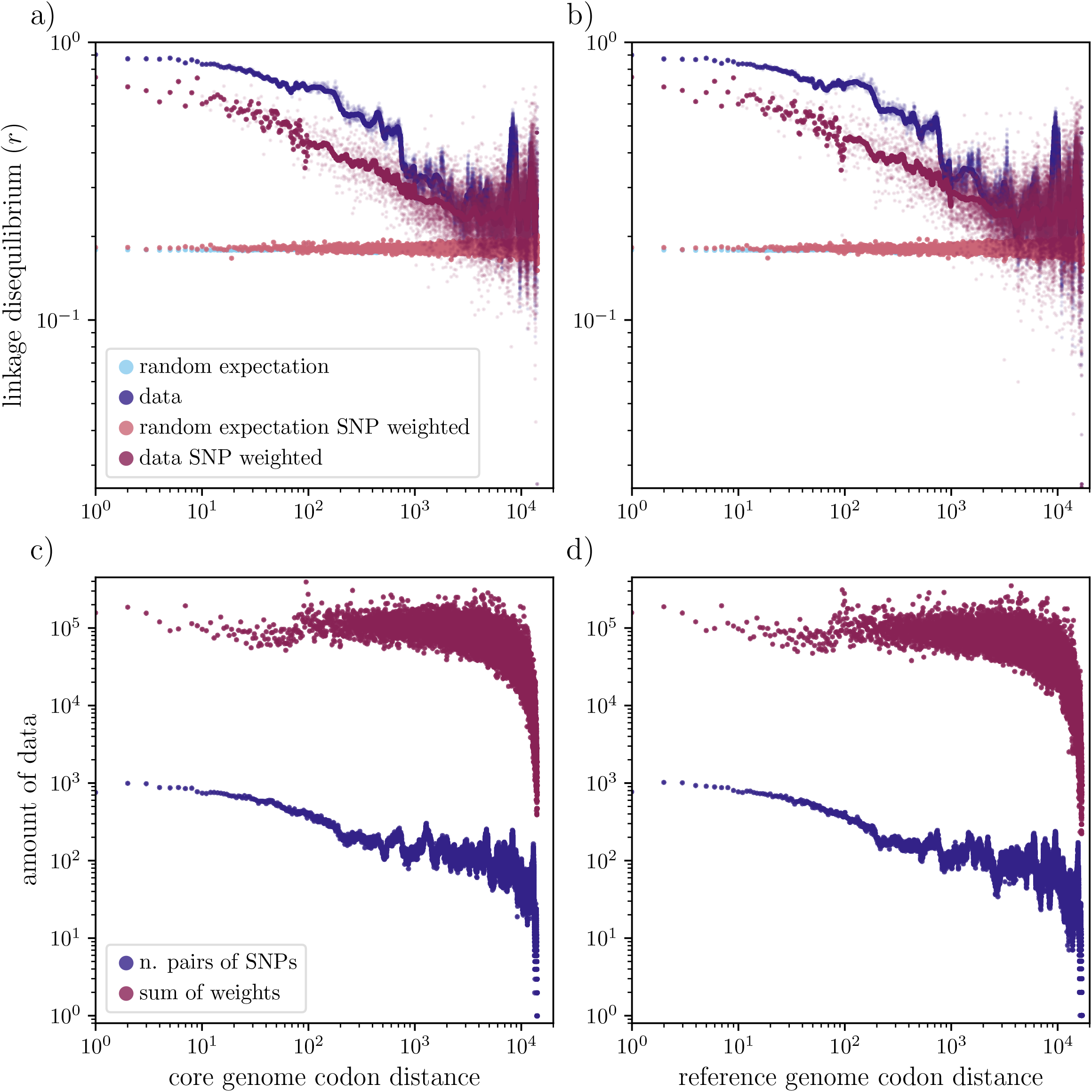
Linkage disequilibrium in group A11. The left hand plots, a) and c), are with respect to codon distance in the core genome, and the right hand plots, b) and d), are with respect to distance within a reference genome, in this case phage MN703405. The figure also shows the effect of weighting the results by SNP density (see SI D.2). The top plots, a)-b), show the linkage disequilibrium with and without the weighting, and the bottom plots, c)-d), show the sum of the weights with the weightings, and the number of data points without weighting. In a)-b), all data points are shown in smaller dots, whereas the larger dots represent a rolling average.

**Figure S7:**
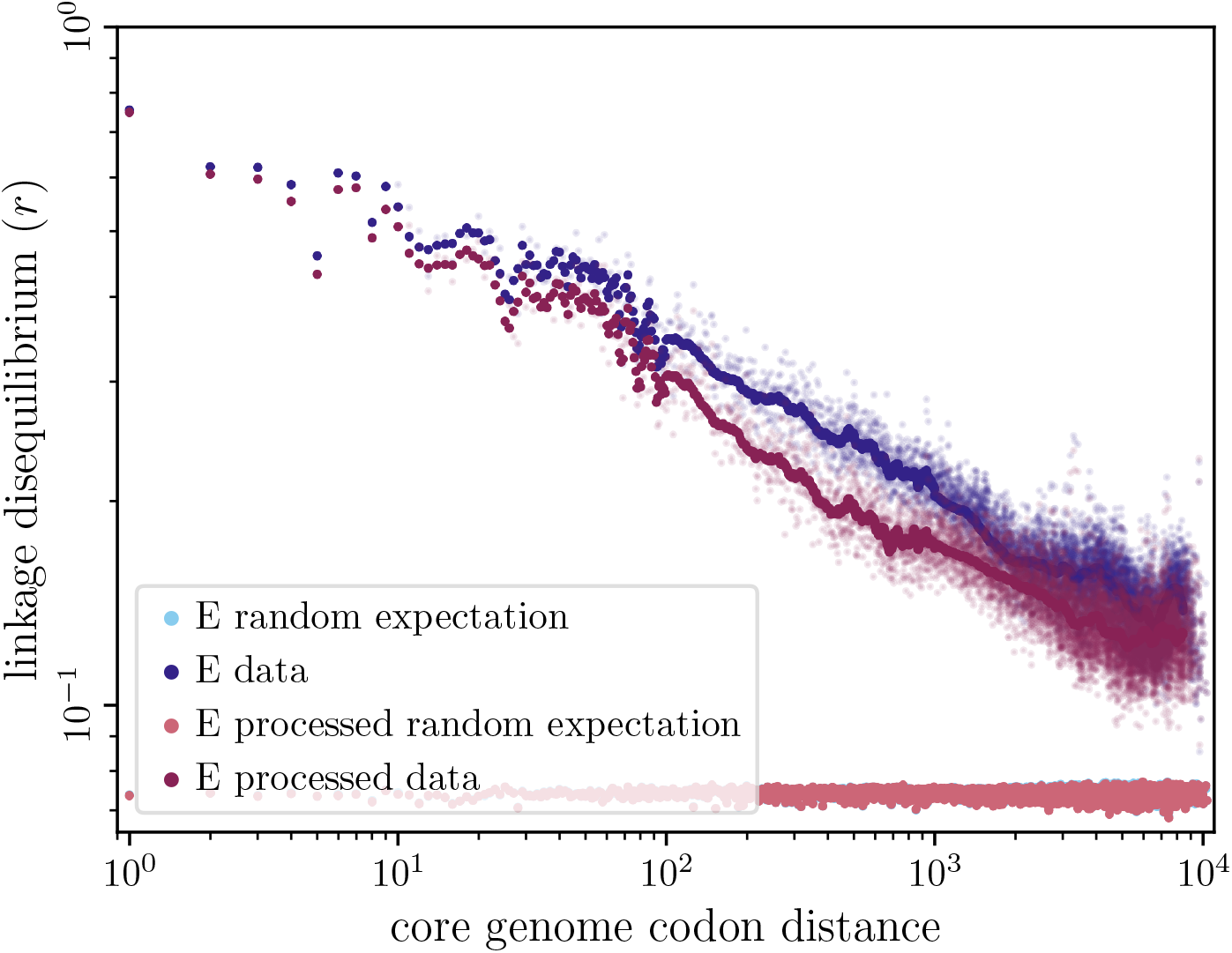
Group E has a small region (approx. 5000 bp) that contains two distinct haplotypes. We recalculated the linkage disequilibrium considering only SNPs that were deviations from the two consensus sequences for each haplotype. The data from this calculation is labeled “E processed” in shades of pink, and the original calculation is shades of blue. The effect of this haplotype region on the linkage disequilibrium is muted since the contribution of high SNP density regions is decreased by the SNP density weighting. However, neglecting the effect still results in a slightly steeper blue curve that decays to a slightly lower asymptotic value.

The small haplotype regions are hard to determine from summary statistics, as the size of the region will affect the contribution to the averages. Moreover, it is worth noting that the manual classifications are not robust, and these characterizations should be thought of as a continuous spectrum, rather than discrete descriptions.

### F SNP compatibility

Inspired by [35], we also investigate the compatibility of sequential SNPs in the core genome alignment. Here, two SNPs are compatible if they pass the 4-allele test. We count the number of SNPs and genomic distance until an incompatible SNP is reached. Failure of the 4-allele test and SNP incompatibility is a signature of recombination.

As the failure of the 4-allele test could be a result of a homoplasy, we also roughly estimate the expected distance between recurrent mutations. Given *N*_*s*_ SNPs out of a core genome alignment of length *N*, we can estimate the mutation number, *K*, as

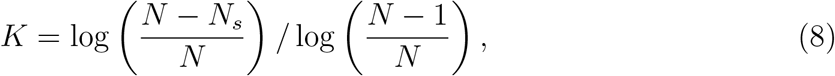

where we have assumed each mutation chooses a site uniformly randomly with probability 1*/N*, and that the SNP sites represent sites that have at least one mutation. Then the expected number of sites that have had more than one mutation, *N*_*r*_, is

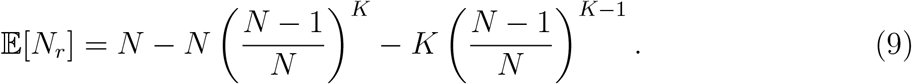

**Figure S8:**
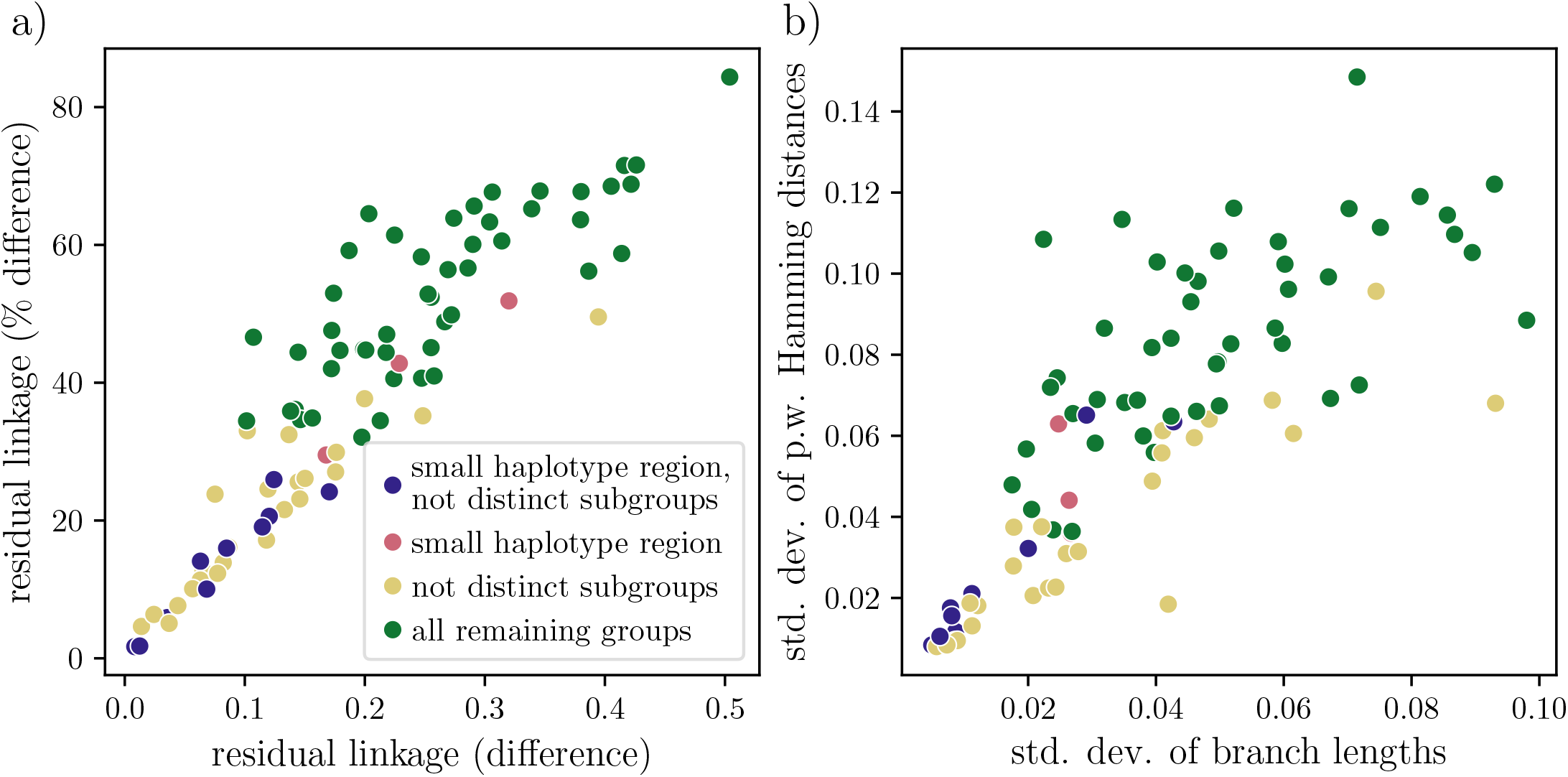
Quantitative characteristics of groups that support manual classifications. a) The amount of residual linkage disequilibrium is a continuous spectrum across all of the groups. High long-range residual linkage disequilibrium is correlated with distinct subgroup structure, but it is not a perfect predictor. b) Properties of the distance matrix of pairwise core Hamming distances also correlate with the manual classifications. The *x*-axis is the standard deviation of the branch lengths of the dendrogram of the distance matrix, and the *y*-axis is the standard deviation of all of the pairwise distances. High standard deviation of either correlates with distinct subgroup structure, but it is again not a perfect predictor.

The estimated distance between recurrent mutations is then simply *N/*𝔼 [*N*_*r*_], and the estimated number of SNPs between recurrent mutations is then simply *N*_*s*_*/*𝔼 [*N*_*r*_]

In all but one of the groups, the estimated distance between recurrent mutations is larger than the mean distance between incompatible SNPs, as shown in S9. This provides evidence that the SNP incompatibility is due to recombination, rather than simply recurrent mutations. Across groups, the unweighted average distance between incompatible SNPs is 66 base pairs.

### G Recombination length estimation

We also wish to estimate the length of recombination events. The linkage decay plots give a rough idea, but to verify those results we looked at divergences between pairs of phages. We considered pairs that were sufficiently diverged such that there could have been a recombination event, but not so diverged that there would be multiple overlapping recombination events. The threshold we used was having a core genome alignment Hamming distance of more than 0.001 but less than 0.02.

We looked at where the two core genome alignments differed as in Fig. S10 a), and plotted the cumulative distribution of the divergences, as in Fig. S10 b). We then chose a maximum of 10 phage pairs per group whose cumulative distribution of SNPs had the highest least squared distance to the diagonal. This was to choose pairs that likely had a single recombination event, rather than mutations scattered across the core genome.

**Figure S9:**
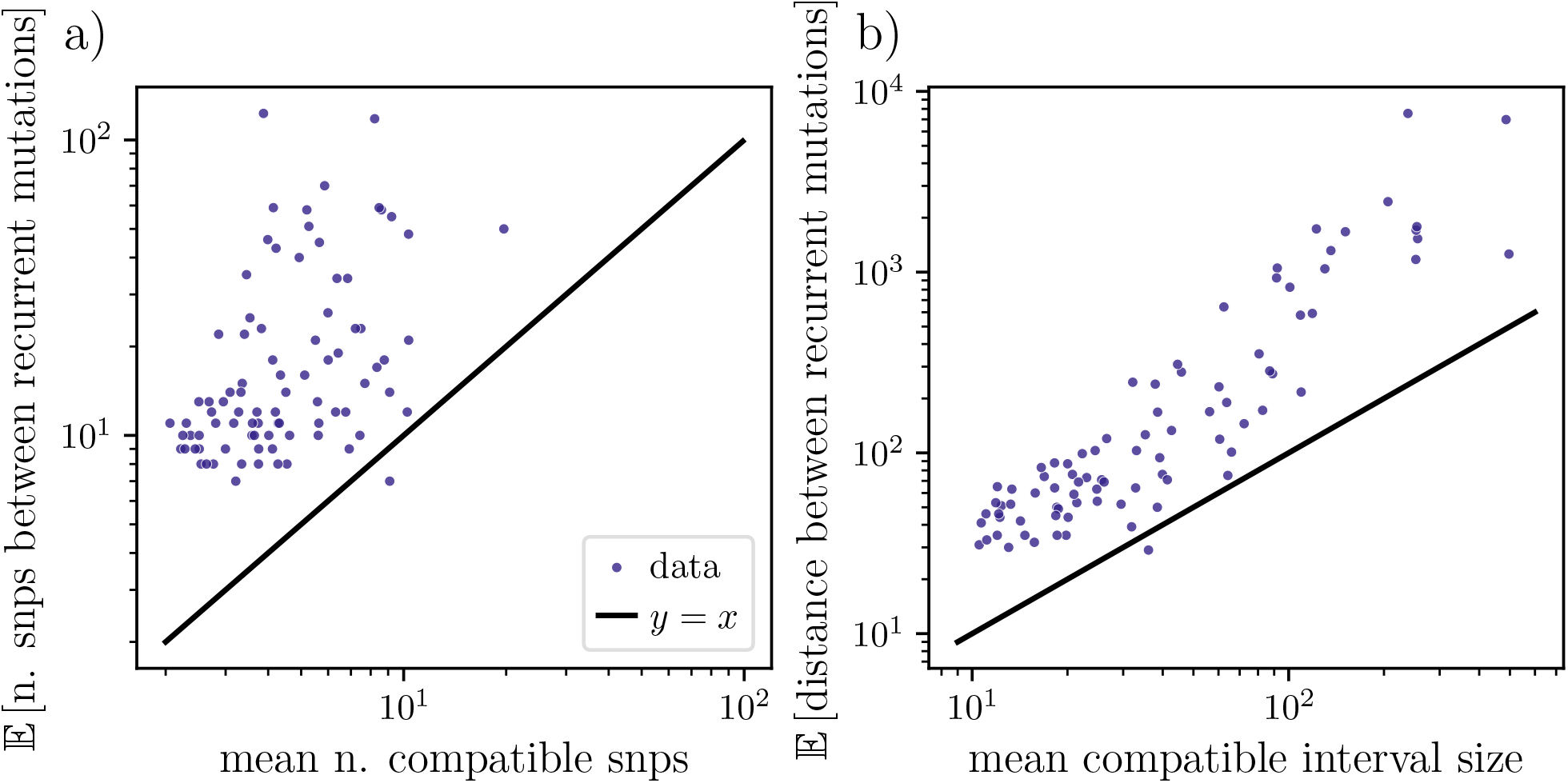
a) The *x*-axis is the mean number of sequential compatible SNPs before an incompatible SNPs is reached. The *y*-axis is the expected number of SNPs between recurrent mutations. b) The *x*-axis is the mean compatible interval size: the base pair distance after which the SNPs become incompatible. The *y*-axis is the expected base pair distance between recurrent mutations.

**Figure S10:**
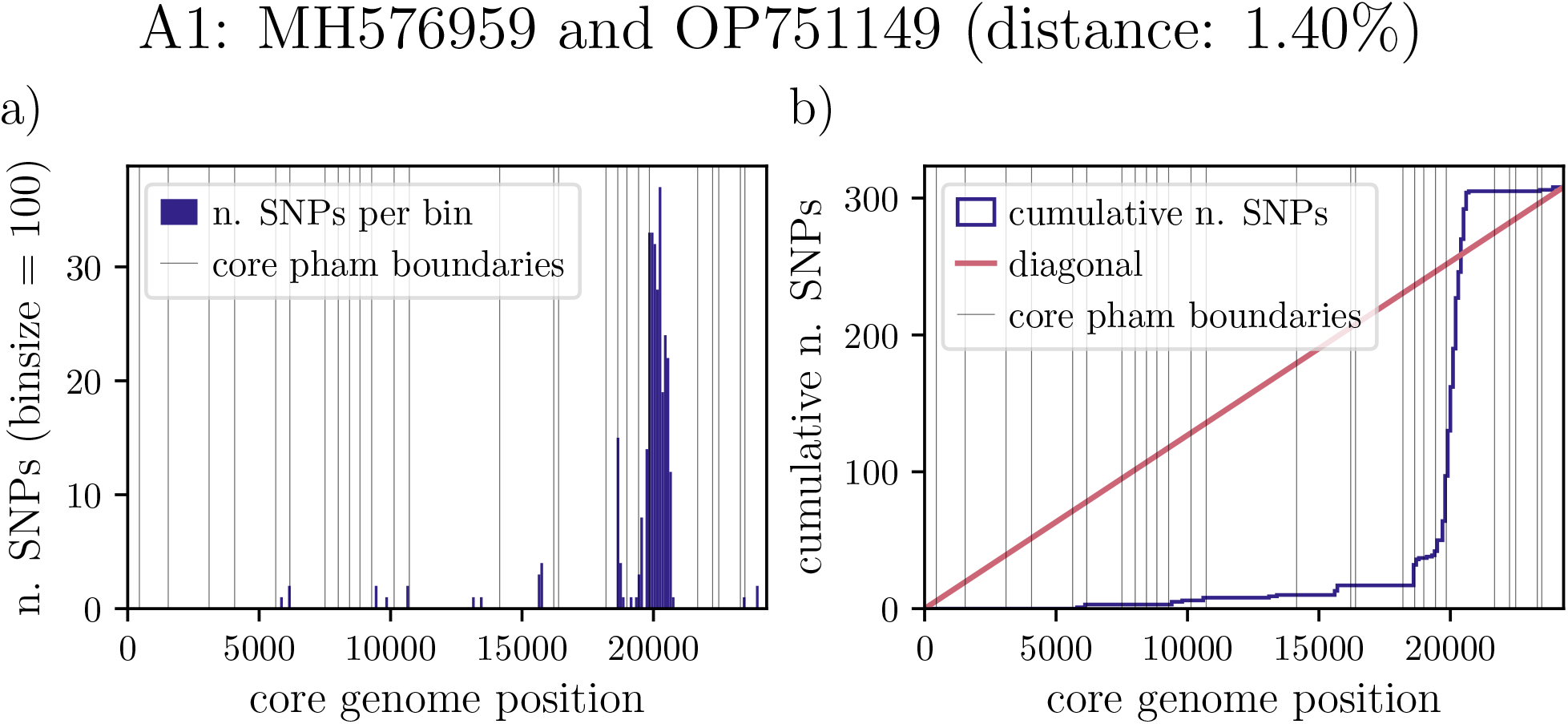
Diagrams representing the divergence between the two core genome alignments of phages MH576959 and OP751149 in group A1. a) shows the number of SNPs across the core genome in each bin of size 100, with the vertical lines representing pham boundaries. b) is the cumulative distribution. The diagonal is plotted to emphasize that cumulative distributions with large deviations from the diagonal are indicators of potential recombination events. In this pair of phages, there is a likely recombination event that occurred just after position 20000, largely within one core pham.

Of these representative phage pairs for each group, we counted the maximum distance over which the divergence was greater than 0.05. In other words, we calculated the maximum length of regions which continuously had at least 5 SNPs in rolling bins of size 100. The majority of bins do not contain at least 5 SNPs, as shown in the distribution in Fig. S10 c).

For each group, we compared the maximum approximate recombination length to the mean length of a core pham, which is shown in Fig. S11. The maximum approximate recombination length is almost always less than the average length of two core phams, and typically less than the length of one core pham. Across groups, the unweighted average of the maximum length of these regions per group is 570 base pairs.

**Figure S11:**
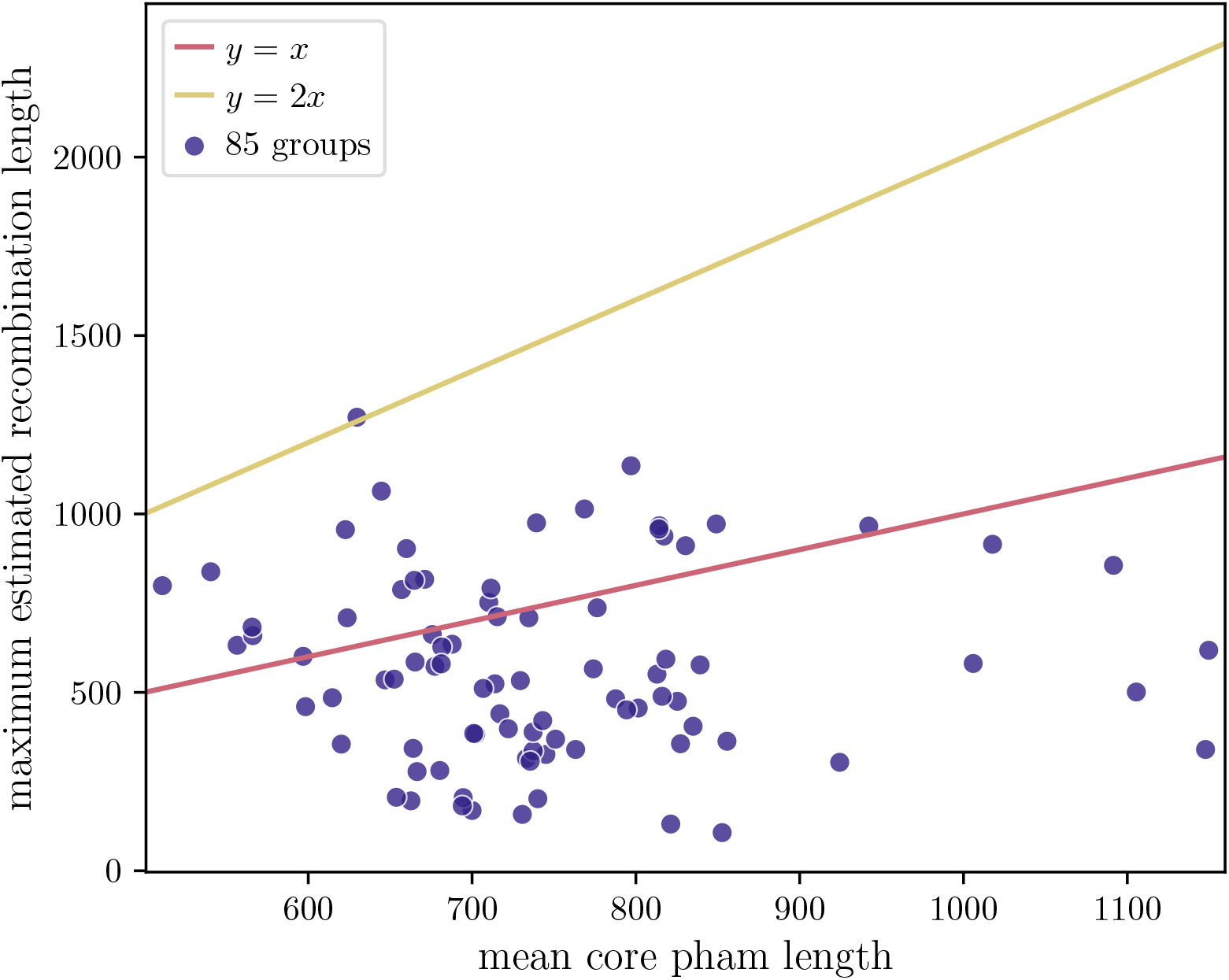
Each dot represents one group. The *x*-axis is the average length of a core pham. The *y*-axis is the maximum estimated recombination length. The pink line is the line *y* = *x*, and the yellow line is the line *y* = 2*x*, displaying that the maximum estimated recombination length is less than or around the average length of one or two core phams. The one group that is missing from the sample did not have any phage pairs that met the divergence thresholds.

